# Delayed localization of A-type lamins to the rupture sites in Hutchinson–Gilford progeria syndrome

**DOI:** 10.1101/2023.09.02.555826

**Authors:** Yohei Kono, Chan-Gi Pack, Takehiko Ichikawa, Arata Komatsubara, Stephen A. Adam, Keisuke Miyazawa, Loïc Rolas, Sussan Nourshargh, Ohad Medalia, Robert D. Goldman, Takeshi Fukuma, Hiroshi Kimura, Takeshi Shimi

**Affiliations:** Cell Biology Center, Institute of Innovative Research, Tokyo Institute of Technology, Yokohama, Japan; Nano Life Science Institute (WPI-NanoLSI), Kanazawa University, Kanazawa, Japan; Faculty of Frontier Engineering, Institute of Science and Engineering, Kanazawa University, Kanazawa, Japan; Convergence Medicine Research Center, Asan Institute for Life Science, Asan Medical Center, Seoul, Korea; Department of Biomedical Engineering, University of Ulsan College of Medicine, Seoul, Korea; School of Life Science and Technology, Tokyo Institute of Technology, Yokohama, Japan; Department of Cell and Developmental Biology, Feinberg School of Medicine, Northwestern University, Chicago, IL; Centre for Microvascular Research, William Harvey Research Institute, Faculty of Medicine and Dentistry, Queen Mary University of London, London, UK; Department of Biochemistry, University of Zurich, Zurich, Switzerland

**Keywords:** Lamin, HGPS, Lamina, Nuclear Envelope Rupture, Nucleus

## Abstract

The nuclear lamina (NL) lines the nuclear envelope (NE) to maintain nuclear structure in metazoan cells. The major NL components, the nuclear lamins contribute to the protection against NE rupture induced by mechanical stress. Lamin A (LA) and a short form of the splicing variant lamin C (LC) are diffused from the nucleoplasm to sites of NE rupture in immortalized mouse embryonic fibroblasts (MEFs). LA localization to the rupture sites is significantly slow and weak compared to LC because of its relatively small pool in the nucleoplasm, but the precise mechanism remains unknown. In this study, we induce NE rupture in wild-type and LA/C-knockout MEFs, and Hutchinson–Gilford Progeria syndrome (HGPS) knock-in MEFs that express progerin, a LA mutant lacking the second proteolytic cleavage site, by laser microirradiation and AFM indentation. The farnesylation at the CaaX motif of unprocessed LA and the inhibition of the second proteolytic cleavage decrease the nucleoplasmic pool and slow the localization to the rupture sites in a long-time window (60-70 min) after the induction of NE rupture. Our data could explain the defective repair of NE rupture in HGPS through the farnesylation at the CaaX motif of unprocessed progerin. In addition, unique segments in LA-specific tail region cooperate with each other to inhibit the rapid accumulation within a short-time window (3 min) that is also observed with LC.

**Significance Statement:** Nuclear lamins are the major components of the nuclear lamina (NL) that lies the nuclear envelope (NE). Lamin A (LA) is slowly localized to sites of nuclear envelope (NE) rupture compared to lamin C (LC). This study reveals that the farnesylation at the CaaX motif of unprocessed LA and the inhibition of the second proteolytic cleavage decrease the nucleoplasmic pool and slow the localization to the rupture sites within a long-time window (60-70 min) after the induction of NE rupture, which could explain the defective repair of NE rupture in Hutchinson–Gilford Progeria syndrome (HGPS). Additionally, unique segments in LA-specific tail region are critical for inhibiting the rapid accumulation within a short-time window (3 min).

## Introduction

In eukaryotic cell nuclei, the nuclear envelope (NE) spatially separates the nuclear genome from the cytoplasm. The barrier function of the NE is due to two phospholipid bilayers of the inner and outer nuclear membranes (INM, ONM). The INM is in contact with chromatin at the nuclear side via INM proteins (1), whereas the ONM is connected with the cytoskeletal system through the linker of nucleoskeleton and cytoskeleton (LINC) complex (2), and is continuous with membranous organelles including the endoplasmic reticulum (ER) (3). The nuclear lamina (NL) is closely apposed to the INM and associates with heterochromatin to support nuclear structure. Nuclear pore complexes (NPCs) fill holes formed by the INM-ONM fusion for nucleo-cytoplasmic transport and interact with euchromatin (4).

The NL structure consists of the nuclear lamins and associated proteins (5). The lamins are type V intermediate filament proteins, which are subdivided into A-type lamins (lamins A [LA] and C [LC]) and B-type lamins (lamins B1 [LB1] and B2 [LB2]) (6, 7). LA and LC are derived from the single gene *LMNA* by alternative splicing (8), whereas LB1 and LB2 are encoded by separate genes *LMNB1* and *LMNB2*, respectively (9-12). Lamin molecules assemble into a filament with a diameter of ∼3.5 nm and these filaments appear to non-randomly lay on top of each other to form meshworks of the NL with a thickness of ∼13.5 nm in immortalized mouse embryonic fibroblasts (MEFs) (13, 14). LA/C and LB1 filaments associate with the outer ring structure of the NPC and are involved in regulating the distribution of the meshworks and NPCs (13-16).

The −CaaX box at the C terminus of unprocessed LA, LB1 and LB2 (pre-LA, pre-LB1 and pre-LB2, respectively) is post-translationally processed in steps, including the attachment of farnesyl group to the cysteine residue of the −CaaX box by a farnesyltransferase (17, 18), proteolytic cleavage of three residues (−aaX) by an aaX endopeptidase (19) and carboxyl methylation of the terminal carboxylic acid group (−COOH) of the C-terminal cysteine residue by a carboxyl methyltransferase (20). The maturation of LB1 and LB2 is completed by these steps. In case of the farnesylated and carboxymethylated pre-LA, an additional 15 C-terminal residues including the farnesylated and carboxymethylated cysteine, are cleaved off by Zinc metallopeptidase STE24 (Zmpste24/FACE1) to produce the mature form (mature-LA) (21, 22).

The NE is locally ruptured during physiological and pathological circumstances. Immediately after NE rupture, NE components are recruited to the rupture sites with the endosomal sorting complex required for transport-III (ESCRT-III) complex and barrier-to-autointegration factor (BAF) (23-25). Nuclear DNA adjacent to the rupture sites is recognized by the DNA sensors cyclic GMP-AMP synthase (cGAS) and its downstream signaling effector STING (23-25). The frequencies of spontaneous and forced NE rupture are significantly increased by the depletion of lamins (26-29). LA/C promotes the accumulation of BAF and cGAS at the rupture sites (30). Numerous mutations that have been found throughout the *LMNA* gene cause a spectrum of human genetic disorders, collectively called laminopathies (31). Laminopathies associated with dilated cardiomyopathy (DCM), muscular dystrophy (MD), familial partial lipodystrophy (FPLD) and Hutchinson–Gilford Progeria syndrome (HGPS) are frequently accompanied by spontaneous NE rupture (32-34). In HGPS, the heterozygous transversion of a nucleotide substitution at position 1824 T>C (G608G) leads to abnormal splicing of the last 150 nucleotides of exon 11, resulting in the production of unprocessed HGPS mutant, progerin (PG) (pre-PG) which lacks 50 amino acids (residues 607–656) from the C terminus of pre-LA (Fig. 1 A). Because these residues contain the second proteolytic cleavage site involved in pre-LA processing, the mature form (mature-PG) remains farnesylated and carboxymethylated, leading to its permanent association with the INM (35, 36).

**Figure 1.**
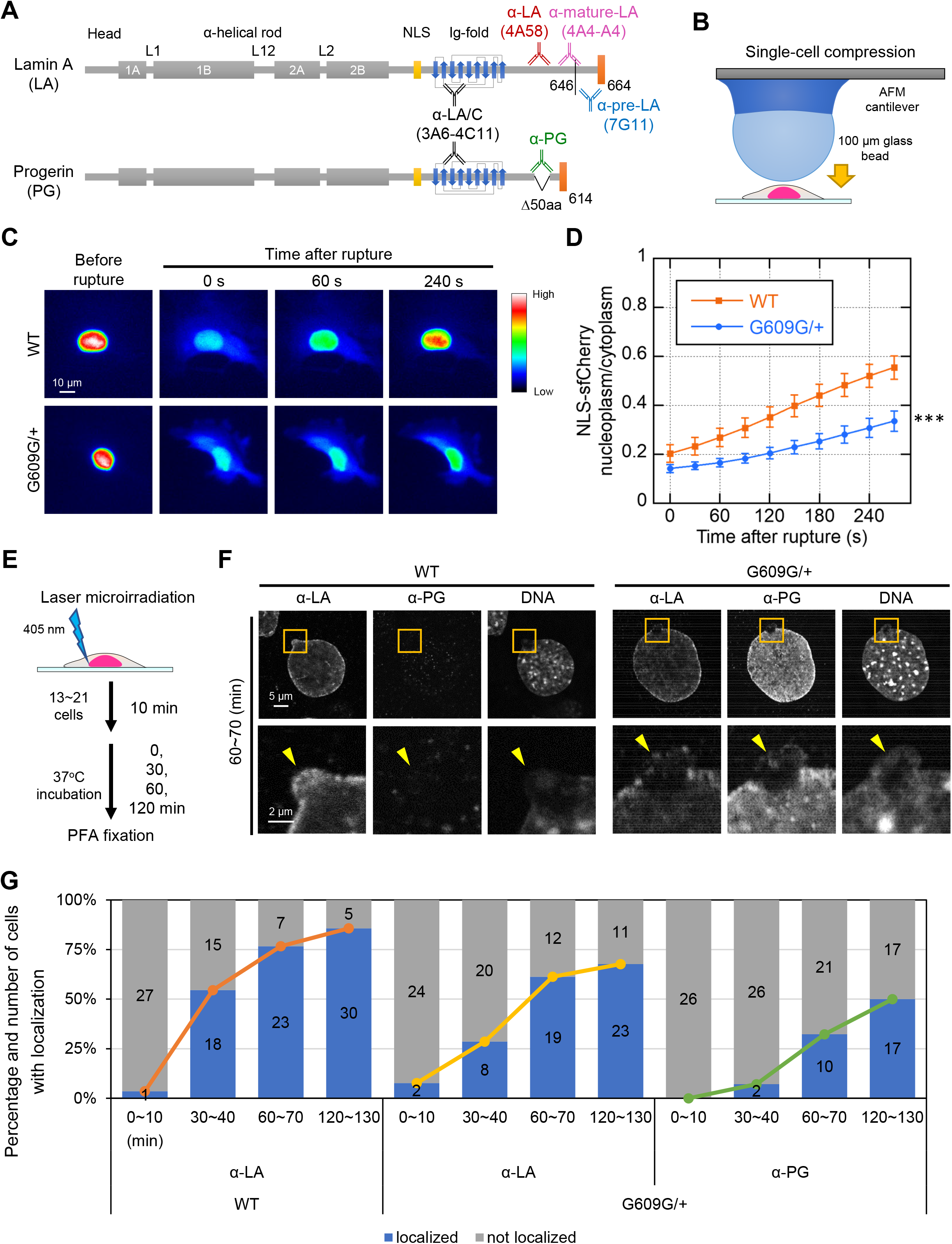
The localization kinetics of LA and PG at the rupture sites in WT and G609G/+ nuclei. **(A)** Protein architecture of LA and PG. The coiled-coil central rod domain (gray), the NLS (yellow), the β-strand comprising the Ig-fold domain (blue), and the CaaX motif (orange) are shown in the diagram. Anti-LA (4A58) and anti-PG do not cross-react with each other. Anti-pre-LA (7G11) recognizes the last amino acids that is proteolytically cleaved off and does not react with PG. Anti-mature-LA (4A4-A4) recognizes mature-LA but not pre-LA. Anti-LA/C (3A6-4C11) recognizes all A-type lamin isoforms. **(B)** NE rupture was induced by AFM indentation using a cantilever with a round tip, as shown in the diagram. **(C** and **D)** Time-lapse images of NLS-sfCherry expressed in WT and G609G/+ cells were acquired with 30 s intervals for 5 min after the induction of NE rupture by AFM indentation. **(C)** The fluorescence intensities in the cells are shown with rainbow color. Bar: 10 μm. **(D)** The nuclear-to-cytoplasmic intensity (N/C) ratios of NLS-sfCherry were measured and normalized to the initial intensities, and plotted to monitor the nuclear reentry in the cells (means ± SEM; n = 20 cells from two independent experiments; ***, P < 0.001 from WT by a mixed effect model). **(E)** NE rupture was induced by 405-nm laser microirradiation with a 2-μm diameter spot at the NE of WT and G609G/+ nuclei. **(F** and **G)** 13 to 21 of nuclei of the cells were laser-microirradiated within 10 min and incubated for 0, 30, 60 and 120 min, followed by fixation with 4% PFA/0.1% Triton-X 100. The fixed cells were immunostained with a combination of the LA and PG antibodies, and the DNA was stained with Hoechst 33342. **(F)** Representative confocal images of nuclei in the cells fixed within 60-70 min after the induction of NE rupture by laser microirradiation. Magnified views of the areas indicated with orange boxes are shown (bottom). The rupture sites are indicated with yellow arrowheads (bottom). Bars: 5 μm (top) and 2μm (bottom). **(G)** Percentiles of the cells with (blue) and without (gray) the localization of LA and PG to the rupture sites. The numbers of analyzed cells from two independent experiments are indicated in the bar charts. The line graphs show the kinetics of the percentage of the cells with localization of LA and PG to the rupture sites.

Previous studies demonstrate that LA/C is recruited to the rupture sites with BAF (30, 37, 38) but LA is significantly slow and weak in localizing to the rupture sites compared to LC (30, 39, 40) because it is less abundant in the nucleoplasm than LC (30, 39, 40). However, a molecular mechanism for retaining nucleoplasmic LA sparse and slowing its localization to the rupture sites still remains to be elucidated. Here, we investigate the role of the C-terminal tail region specific for LA in this mechanism. We previously showed that mEmerald-fused LA accumulated at the rupture sites only when it was overexpressed at a high level (30), and the accumulation kinetics appeared to be artifactual because it was not confirmed with endogenous LA detected by immunofluorescence (30). Therefore, we establish the ectopic expression system for live-cell imaging and fluorescence correlation spectroscopy (FCS) to express mEmerald-fused LA, alternate forms of LA, and LC at a low level with the endogenous promotor in wild type (WT), HGPS and LA/C-knockout (KO) MEFs. Our data reveal the significance of the LA-specific tail region for retaining its small pool in the nucleoplasm and slowing its localization to the rupture sites.

## Results

### PG is not only slow in localizing to the rupture sites but also slows LA localization

To investigate the involvement of the pre-LA processing in retaining small pool of LA in the nucleoplasm and slowing its localization to the rupture sites, we used immortalized WT MEFs and heterozygous knock-in MEFs with PG expression from a mutant allele of the *Lmna* gene carrying the c.1827C>T; p.Gly609Gly mutation (*Lmna^G609G/+^*, G609G/+). This mutation is equivalent to the HGPS mutation in the human *LMNA* gene (41, 42). These cells stably expressed super-folder Cherry harboring two NLSs derived from SV40 large T antigen and c-Myc (NLS-sfCherry) as a NE rupture marker (30). Because spontaneous NE rupture is frequently observed in skin fibroblasts derived from a HGPS patient and mouse smooth muscle cells (SMCs) expressing PG (32, 34), we first examined if G609G/+ MEFs also exhibit misshapen nuclei, the leakage of NLS-sfCherry to the cytoplasm, the mis-localization or decreased expression of LB1, and the local accumulation of cGAS, which are all indicative of spontaneous NE rupture. WT and G609G/+ cells were stained with Hoechst 33342 for DNA and specific antibodies directed against LB1 and cGAS (Fig. S1 A). Unlike the HGPS patient cells and PG-expressing mouse SMCs, G609G/+ cells did not show these phenotypes compared to WT cells (Table S1), indicating that the HGPS mutation is not accompanied by spontaneous NE rupture in these cells. Next, to examine the effect of this mutation on recovery from forced NE rupture, single-cell compression by AFM indentation was applied to these cells, and the nuclear reentry of NLS-sfCherry was observed by time-lapse imaging before and after the induction of the rupture (Fig. 1 B) (43). Interestingly, G609G/+ cells showed a significantly slow reentry compared to the WT cells which had a ∼50% recovery from the original fluorescence intensity ∼4 min after the induction of NE rupture (Fig. 1 C and D), indicating that PG expression slows the recovery from forced NE rupture.

To determine the localization kinetics of LA and PG at the rupture sites within a relatively long-time window, WT and G609G/+ cells were fixed at 0-10, 30-40, 60-70 and 120-130 min after the induction of NE rupture by 405-nm laser microirradiation (Fig. 1 E) (30), and then stained with Hoechst 33342 for DNA and the specific antibodies directed against LA and PG which do not cross-react to each other (Fig. 1 A). At 0-10, 30-40, 60-70 and 120-130 min after the rupture, LA was localized to the rupture sites in ∼4%, ∼55%, ∼77% and ∼86% of WT nuclei and ∼8%, ∼29%, ∼61% and ∼68% of G609G/+ nuclei, respectively (Fig. 1 F, G and Fig. S1 B), whereas PG was slower in localizing to the rupture sites of G609G/+ nuclei than LA, showing ∼0%, ∼7%, ∼32% and ∼50%, respectively (Fig. 1 F, G and Fig. S1 B). These results suggest that PG expression decelerates the localization kinetics of LA at the later time points (60 and 120 min). In addition, both LA and PG are likely to be recruited to the rupture sites with BAF within a relatively long-time window according to our results from BAF knockdown (KD) experiments with short hairpin RNAs (shRNAs) (Fig. S1 C-E). WT cells expressing a shRNA for the control (Scramble) or two shRNAs (shBAF #1 and #2) were fixed at 60-70 min after the induction of NE rupture by laser microirradiation, and then LA localization was determined by immunofluorescence. As expected, LA was localized to the rupture sites in ∼61%, 10% and ∼16% of the control and BAF-KD#1 and #2 nuclei, respectively (Fig. S1 D and E), indicating that BAF is required for LA localization to the rupture sites. These results are similar to a previous finding to indicate that BAF rapidly recruits overexpressed GFP-LA to the rupture sites (37).

### PG slows LC accumulation to the rupture sites

Because the accumulation of LC to the rupture sites was much earlier (within 3 min) than the localization of LA or PG (30-120 min) (30, 39, 40) (Fig. 1 F and G) and LC depletion accelerates the cytoplasmic leakage of NLS-sfCherry at the early time points (within 10 min) (30), LC might be directly linked to the delay in retrieving NLS-sfCherry from the cytoplasm of G609G/+ cells within 10 min after the induction of forced NE rupture (Fig. 1 C and D). Therefore, PG expression could slow LC accumulation to the rupture sites. To test this idea, endogenous LA, LC and PG were directly labeled with a LA/C-specific genetically encoded probe, designed ankyrin repeat protein (DARPin)-LaA_6 (44), fused with super-folder GFP (sfGFP), sfGFP-DARPin-LA6 in WT and G609G/+ cells expressing NLS-sfCherry (30) (Fig. S2 A). Then, live-cell imaging was carried out before and after the induction of NE rupture by laser microirradiation. Because endogenous LA and PG did not localize to the rupture sites within 3 min after the induction of NE rupture (30) (Fig. 1 G), sfGFP-DARPin-LA6 accumulating at the rupture sites only represented endogenous LC within such a short-time window. Immediately after the induction of NE rupture, the accumulation of sfGFP-DARPin-LA6 was significantly weaker in G609G/+ nuclei compared to WT nuclei (Fig. S2 B and C), indicating that PG expression attenuates LC accumulation.

### PG slows the diffusion of LC in the nucleoplasm

Because of the slow recruitment of LC from the nucleoplasm to the rupture sites in G609G/+ nuclei (Fig. S2 B and C), the diffusion fractions and coefficients in the nucleoplasm could be decreased in G609G/+ nuclei compared to WT nuclei. To determine them, mEmerald-LC was stably expressed at a low level from the *Rosa26* locus under the endogenous *Lmna* promotor in WT and G609G/+ cells, and fluorescence correlation spectroscopy (FCS) was performed using these cells (Fig. S3 A-D). The auto-correlation curve obtained by the signal detection was fitted with fast and slow components (15, 45, 46). Interestingly, the diffusion coefficients of fast components were significantly high in WT nuclei compared to G609G/+ nuclei (Fig. S3 D). This result indicates that PG expression significantly decreases the diffusion coefficient of fast component despite the increased fraction but this degree of difference is insufficient for explaining the weak accumulation of sfGFP-DARPin-LA6 at the rupture sites in G609G/+ nuclei compared to WT nuclei.

### The nucleoplasmic pools of LA and PG are increased by the inhibition of farnesylation

When the CaaX motifs of pre-LA and pre-PG are farnesylated (18, 36, 47), they are retained at the NE through the hydrophobic interaction with the lipid bilayer of the INM (47, 48). Therefore, the inhibition of farnesylation with FTI could increase their nucleoplasmic pools, leading to their fast localization to the rupture sites. To test this possibility, WT and G609G/+ cells were treated with DMSO for the control or farnesyltransferase inhibitor (FTI), 3.2 µM lonafarnib for 48 h, and the expression levels and localization of LA and PG were determined by immunoblotting using the specific antibodies directed against LA/C, and by immunofluorescence using the LA and PG antibodies respectively, (Fig. 2 A). For immunoblotting, pre-LA, mature-LA, pre-PG, mature-PG, and LC were detected with the antibodies directed against LA/C (for all) and PG (for pre- and mature-PG). For immunofluorescence, the LA antibody (for pre- and mature-LA) and the PG antibody (for pre- and mature-PG) were used. As expected, the levels of pre-LA and pre-PG were increased in FTI-treated cells compared to the control cells as shown as the upper and lower band of LA and PG (Fig. 2 A). Based on ratios of the average fluorescence intensity in the nucleoplasmic pool relative to that of the NL (NP/NL ratios), LA and PG were significantly abundant in the nucleoplasm of FTI-treated nuclei compared to the control nuclei (Fig. 2 B and C), indicating that LA and PG are increased in the nucleoplasmic pool by FTI treatment. Next, these cells were fixed within 10 min and at 60-70 min after the induction of NE rupture by laser microirradiation, and the localization of LA and PG to the rupture sites was detected by immunofluorescence. LA was localized to the rupture sites within 10 min in ∼7% and ∼7% of WT and G609G/+ nuclei for the control and ∼18% and ∼17% of WT and G609G/+ nuclei treated with FTI, respectively, and PG in 0% and ∼37% of the control and FTI-treated G609G/+ nuclei, respectively (Fig. S4). At 60-70 min, LA localization to the rupture sites was observed in ∼59% and ∼42% of the control nuclei and ∼91% and ∼74% of FTI-treated WT and G609G/+ nuclei, respectively, and PG localization in ∼18% and ∼81% of the control and FTI-treated G609G/+ nuclei, respectively (Fig. 2 D). These results indicate that the inhibition of farnesylation promotes the localization of LA and PG to the rupture sites.

**Figure 2.**
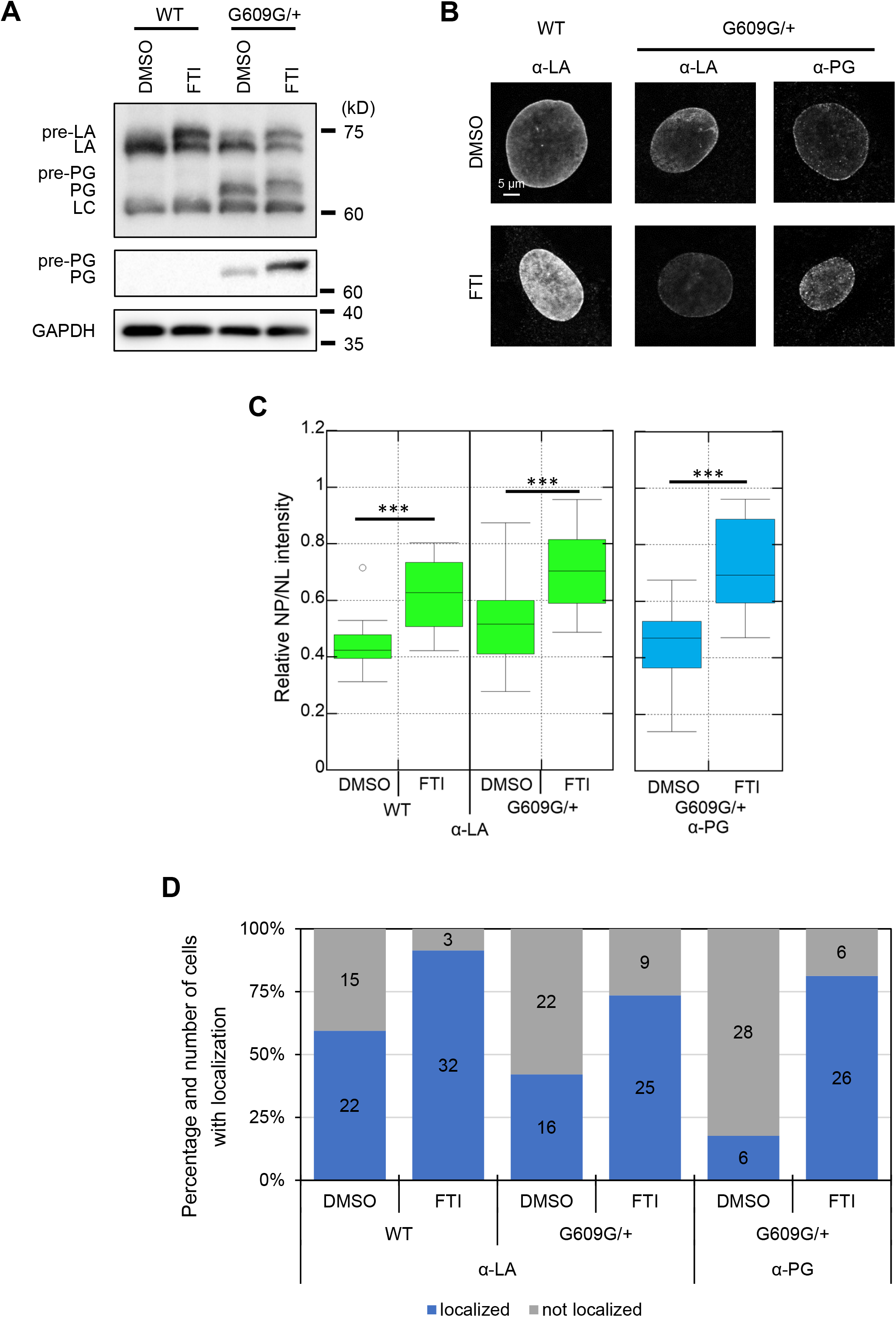
Localization of LA and PG to the rupture sites in DMSO or FTI-treated WT and G609G/+ nuclei. **(A)** Whole cell lysates from WT and G609G/+ cells treated with 0.1% DMSO or 3.2 μM lonafarnib (FTI) were probed with anti-LA/C (3A6-4C11), anti-PG, and anti-GAPDH (as loading control) for immunobloting. Positions of the size standards are shown on the right. **(B** and **C)** The indicated cells were immunostained with the LA (4A58) and PG antibodies. **(B)** Representative microscopic images. **(C)** Ratios of the average fluorescence intensity in the nucleoplasmic pool relative to that of the NL (NP/NL ratios) of LA and PG were measured based on immunofluorescence (n = 20 cells from two independent experiments; ***, P < 0.001 by Welch’s t-tests). **(D)** 15 to 19 of nuclei in the cells were laser-microirradiated and incubated for 60 min, followed by fixation with 4% PFA/0.1% Triton-X 100. The fixed cells were immunostained with a combination of the LA and PG antibodies. Percentiles of the cells with (blue) and without (gray) the localization of LA and PG at the rupture sites. The numbers of analyzed cells from two independent experiments are indicated in the bar charts.

### PG becomes diffusible in the nucleoplasm by FTI treatment

Because of the increases in nucleoplasmic pool of LA and PG by FTI treatment (Fig. 2 B and C), the non-farnesylated forms could be highly mobile in the nucleoplasm of FTI-treated cells. To test this idea, the diffusion coefficients of mEmerald-LA and PG were measured by FCS (Fig. 3 A). For the FCS measurements, mEmerald-LA and PG were stably expressed at a low level by the same system as described above with mEmerald-LC (Fig. S3 A and B), followed by the treatment with DMSO for the control or FTI (Fig. S5). The diffusion fractions of mEmerald-LA in the control and FTI-treated WT and G609G/+ nuclei were all similar to each other (Fig. 3 B and C) but the diffusion coefficient of fast component was slightly but significantly small in G609G/+ nuclei compared to WT nuclei whereas that of slow component remained unchanged (Fig. 3 C). These results show that PG expression slows the diffusion speed of mEmerald-LA. On the other hand, the diffusion signals of mEmerald-PG were barely detectable in the nucleoplasm of WT and G609G/+ nuclei for the control. Instead, the signal decays caused by photobleaching were observed during the FCS measurements (Fig. 3 D), strongly suggesting that these signals originate from the NL where mEmerald-PG are immobile (15, 53). When WT and G609G/+ cells were treated with FTI, the diffusion signals were detected by FCS as expected (Fig. 3 D and E), indicating that PG becomes diffusible in the nucleoplasm by FTI treatment. Therefore, the nucleoplasmic pool is the likely source of PG localized to the rupture sites (Fig. 2 C and D).

**Figure 3.**
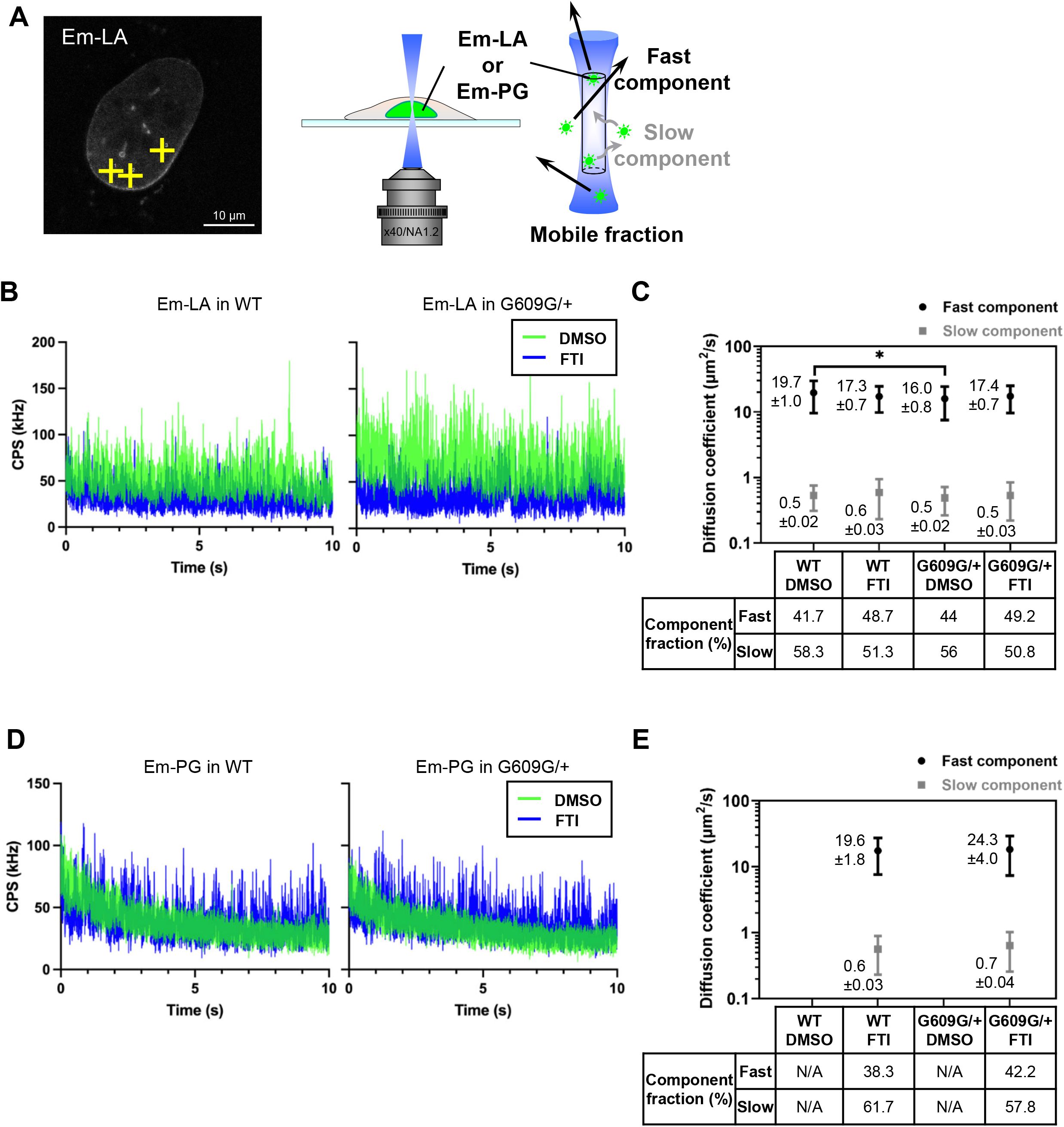
The diffusion kinetics of LA and PG in the nucleoplasm of DMSO or FTI-treated WT and G609G/+ nuclei. The diffusion fractions of mEmerald-LA (Em-LA) and mEmerald-PG (Em-PG) in DMSO- or FTI-treated WT and G609G/+ nuclei were measured by FCS. **(A)** A representative confocal image of a WT nucleus with Em-LA before FCS measurements. The yellow crosses indicate the points measured by FCS (left). Bar: 10 µm. Em-LA or Em-PG molecules move in or out of the confocal volume (white-out cylinder region in blue) at different speeds, as shown in the diagram (right). **(B)** Representative fluorescence fluctuations of Em-LA in DMSO-(green line) and FTI-treated (blue line) of WT and G609G/+ nuclei are plotted by count per second (CPS). **(C)** Diffusion coefficients of fast component (plotted in black) and slow component (plotted in gray) for Em-LA in the cells. **(D)** Representative fluorescence fluctuations of Em-PG in DMSO-(green line) and FTI-treated (blue line) of WT and G609G/+ nuclei are plotted by CPS. **(E)** Diffusion coefficients of fast component (plotted in black) and slow component (plotted in gray) for Em-PG in the cells. **(C** and **E)** Mean ± SEM are indicated on the left to plots; n = 10 cells from two independent experiments; *, P < 0.05 by a Games–Howell test. Percentiles of these component fractions are indicated at the bottom of the graph. N/A, not applicable because of photobleaching.

### The second proteolytic cleavage is required for the localization of LA to the rupture sites

The localization of PG to the rupture sites was significantly slow compared to LA (Fig. 1 G), and the permanent association of PG with the INM due to the absence of the second proteolytic cleavage site could decelerate the localization kinetics (35, 36). To check this possibility, Zmpste24, which endoproteolytically processes both the CaaX motif and the second cleavage site (54) was knocked down by shRNAs in WT MEFs, and then the expression levels and localization of pre- and mature-LA were determined in cells expressing shRNAs for the control or Zmpste24 KD by immunoblotting and immunofluorescence, respectively (Fig. S6 A). A specific antibody directed against pre-LA used for immunoblotting and immunofluorescence reacts with pre-LA but not mature-LA (Fig. 1 A). The C-terminal residues of pre-LA including the CaaX motif remained partially uncleaved by Zmpste24 KD, resulting in the significant increase and decrease in expression level of pre- and mature-LA, as shown as the upper and lower band, respectively (Fig. S6 A). These results are consistent with the previous findings to indicate that Zmpste24 KD inhibits the second cleavage (55). LA and pre-LA were predominantly localized to the NL, and the NP/NL ratios remained unchanged in Zmpste24-KD cells compared to the control cells (Fig. S6 B and C). These cells were fixed at 60-70 min after the induction of NE rupture by laser microirradiation, and then the localization of LA and pre-LA to the rupture sites was determined by immunofluorescence. LA was localized at the rupture sites in ∼75% of the control cells whereas LA and pre-LA were completely absent from the rupture sites in Zmpste24-KD cells (Fig. S6 D and E), indicating that the second cleavage is required for their localization to the rupture sites.

### Inhibition of farnesylation at the CaaX motif is sufficient for the localization of LA to the rupture sites without the second proteolytic cleavage

Zmpste24 depletion significantly increased the pre-LA expression level (56) (Fig. S6 A and B), which is accompanied by the farnesylation at the CaaX motif (49). Therefore, the inhibition of the farnesylation with FTI could release the pre-LA from the NE, resulting in the fast localization of pre-LA to the rupture sites. To test this idea, the control (Scramble) and Zmpste24-KD cells were treated with DMSO or FTI as before, and the expression levels and localization of pre- and mature-LA were determined by immunoblotting and immunofluorescence, respectively (Fig. 4 A and B). As expected, Zmpste24 KD increased and decreased the pre- and mature-LA levels, respectively in DMSO- and FTI-treated cells (Fig. 4 A). FTI treatment also had the similar effect in the control cells but the synergetic effect with Zmpste24 KD was very subtle (Fig. 4 A). To analyze the localization of pre- and mature-LA in those cells by immunofluorescence, specific antibodies directed against them that do not cross-react with each other were used. In the control cells treated with DMSO, pre-LA was under the detection limit whereas mature-LA was predominantly localized to the NL (Fig. 4 B-D). When the control cells were treated with FTI, pre-LA became clearly visible in the NL and mature-LA was less prominent there instead (Fig. 4 B-D). In Zmpste24-KD cells treated with DMSO, pre-LA was localized to the NL and mature-LA was undetectable (Fig. 4 B-D). FTI treatment decreased pre-LA localization to the NL up to the level of the control cells treated with FTI and mature-LA was undetectable (Fig. 4 B-D). These results were consistent with the known effect of FTI treatment, which inhibit the farnesylation at the CaaX motif and the following processes including the second cleavage, leading to release the pre-LA from the NE (50). As FTI treatment increases nucleoplasmic LA levels regardless of the second cleavage, we hypothesize that LA localizes to the rupture site more efficiently in FTI. The localization of pre- and mature-LA to the rupture sites was determined in these cells by immunofluorescence at 60-70 min after the induction of NE rupture by laser microirradiation. Pre-LA was localized to the rupture sites in ∼97%, ∼12% and ∼77% of FTI-treated control, DMSO- and FTI-treated Zmpste24-KD nuclei, respectively (Fig. 4 E). This indicates that pre-LA without farnesylation present in the nucleoplasm effectively localizes to the rupture sites, supporting the hypothesis. However, the localization of mature-LA was slightly decreased in the control cells by FTI treatment (Fig. 4 E), despite FTI treatment increases the localization of pre- and mature-LA using antibody directed against both (Fig. 2 D), probably because its nucleoplasmic level was not as high as pre-LA (Fig. 2 C, 4 C and D).

**Figure 4.**
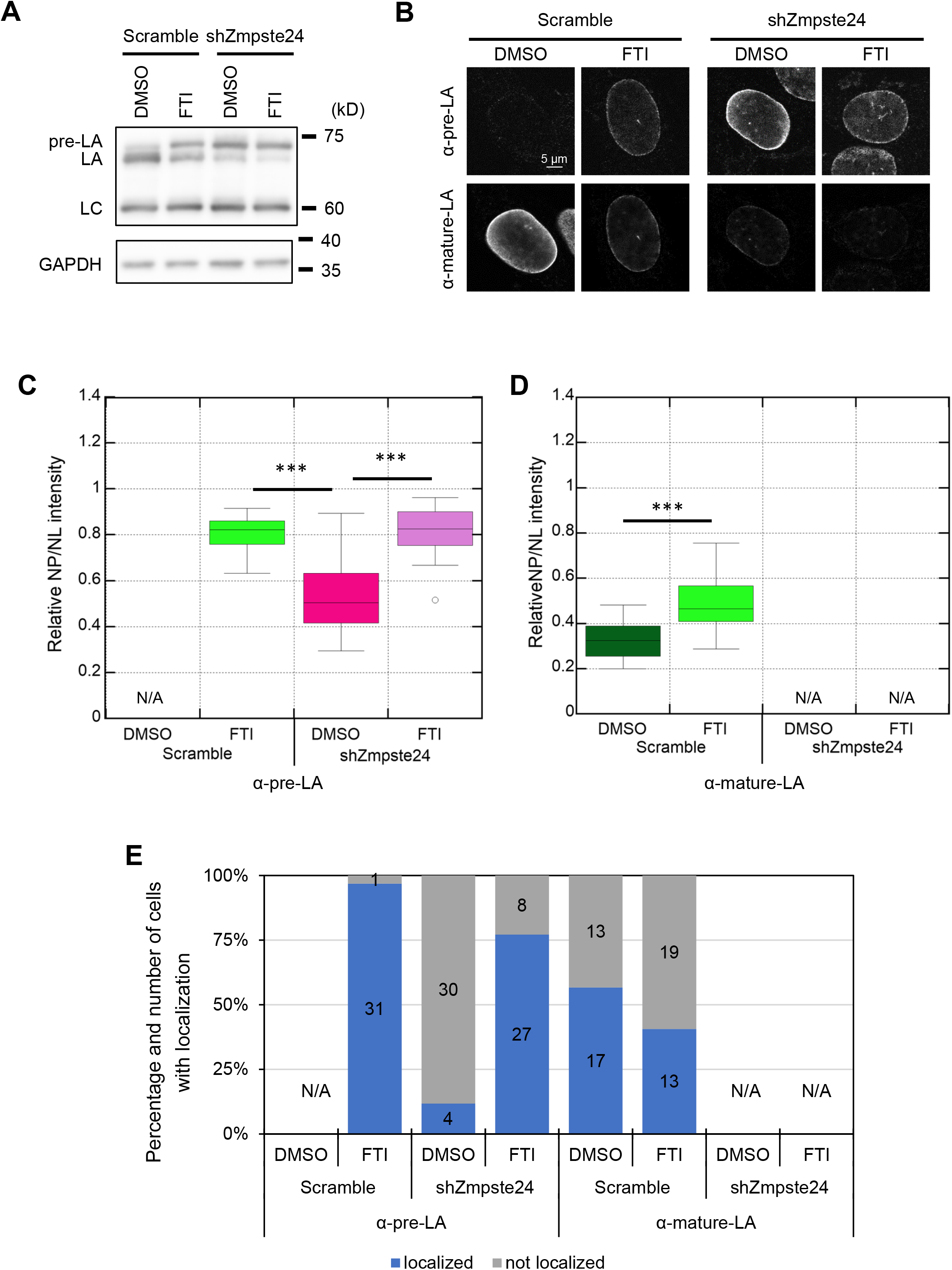
The localization of pre- and mature-LA to the rupture sites in the control and Zmpste24-KD WT MEFs treated with DMSO or FTI. **(A)** Whole cell lysates from the control and Zmpste24-KD cells treated with DMSO or FTI were probed with anti-LA/C (3A6-4C11) and anti-GAPDH (as loading control). Positions of the size standards are shown on the right. **(B)** The control and shZmpste24 cells treated with DMSO or FTI were immunostained with a combination of anti-pre-LA (7G11) and anti-mature-LA (4A4-A4). Bar: 5 μm. **(C** and **D)** NP/NL ratios of pre-LA (C) and mature-LA (D) in the cells were measured based on immunofluorescence (n = 20 cells from two independent experiments; ***, P < 0.001 by Games–Howell tests for pre-LA and by Welch’s t-test for mature-LA). **(E)** 15 to 18 of nuclei in the cells were laser-microirradiated within 10 min and incubated for 60 min, followed by fixation with 4% PFA/0.1% Triton-X 100. The fixed cells were immunostained with a combination of the mature-LA and pre-LA antibodies. Percentiles of the indicated cells with (blue) and without (gray) the localization of LA and pre-LA to NE rupture sites. The numbers of analyzed cells from two independent experiments are indicated in the bar charts. **(C**-**E)** N/A, not applicable due to below detection limit.

### LA differs from LC in localization to the rupture sites through two specific sequences in the tail region

The inhibition of farnesylation at the CaaX motif of PG significantly increased the nucleoplasmic pool and promoted the localization to the rupture sites within 10 min after the induction of NE rupture (Fig. S4). Interestingly, PG was not localized to the rupture sites within the first half of the time window (0-5 min) whereas LC rapidly accumulated (30) (Fig. S2). Therefore, we confirmed the effect of this inhibition on their NP/NL ratios and the accumulation kinetics at the rupture sites within the short-time window (3 min) by live-cell imaging using mEmerald-fused LA-full length (FL), LC, PG and the non-farnesylated mutant of PG, PG-CSM that lacks the isoleucine within the CaaX motif were expressed in LA/C-KO MEFs (Fig. 5 A). As expected, the NP/NL ratio of mEmerald-PG was similar to that of mEmerald-LA-FL, whereas that of mEmerald-PG-CSM was significantly high but not to the extent of mEmerald-LC (Fig. 5 B and C). mEmerald-PG and mEmerald-PG-CSM did not accumulate at the rupture sites (Fig. 5 D), indicating that the inhibition of the farnesylation is not sufficient for exhibiting the rapid accumulation that was similar to that of mEmerald-LC. These results strongly suggest that the tail regions specific for LA and PG are involved in inhibiting the rapid accumulation. In human and mice, LC harbors identical six amino acids whereas LA has 98 and 97 amino acids, following the residues 1-566 and 1-568 shared between LA and LC, respectively (Fig. S7 A). Because this LC-specific amino acids do not contribute to the rapid accumulation at the rupture sites (30), mEmerald-fusion proteins of LA truncation mutants, LA-S575X, LA-A601X, LA-S617X, LA-F627X, LA-T644X and LA-L648X were tested for their NP/NL ratios and the accumulation kinetics within 3 min (Fig. 5 A). Among these mutants, mEmerald-LA-S575X has the highest NP/NL ratio (Fig. 5 B and C), and only this mutant rapidly accumulated within 3 min (Fig. 5 D and S7 B), indicating that the residues 575-600 is involved in inhibiting the rapid accumulation. Because the deletion mutant, mEmerald-LAΔ26 did not accumulate at the rupture sites (Fig. 5 D), further truncation was performed at the C-terminal side of mEmerald-LAΔ26 to produce mEmerald-LAΔ26-L627X, mEmerald-LAΔ26-L644X and mEmerald-LAΔ26-L648X (Fig. 5 A). The NP/NL ratio remained unchanged by these truncations compared to mEmerald-LAΔ26 (Fig. S7 C and D) but only mEmerald-LAΔ26-L627X rapidly accumulated (Fig. 5 E and S7 E). These results show that the residues 627-643 assists LAΔ26 to inhibit the rapid accumulation without decreasing the nucleoplasmic pool. We therefore designated the residues 575-600 as LA-characteristic sequence-1 (LACS1) and the residues 627-643 as LACS2.

**Figure 5.**
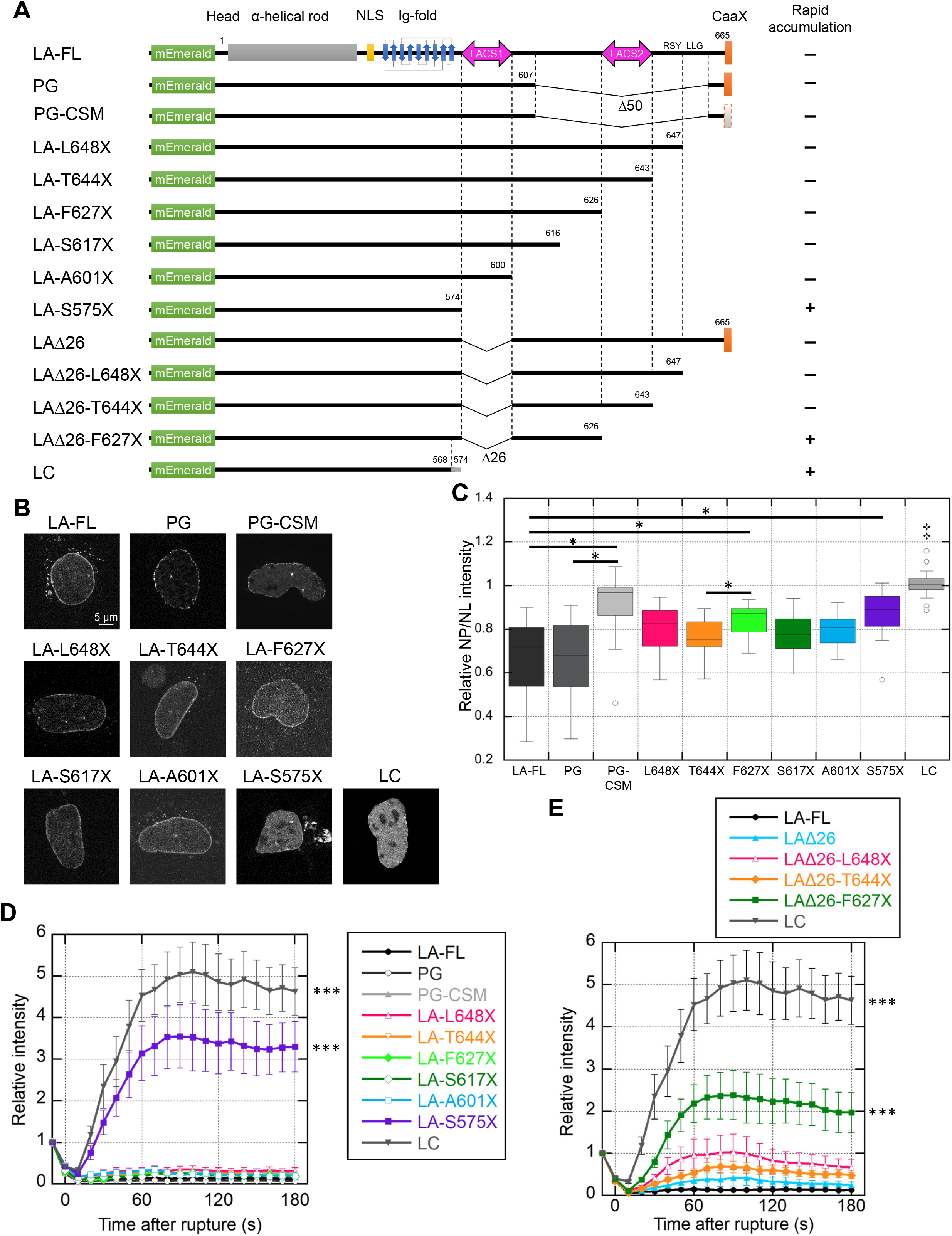
Rapid accumulation of LA truncation and PG deletion mutants at the rupture sites in LA/C-KO nuclei. **(A)** Architecture of mEmerald-fused LA-full-length (FL), PG, PG-CSM, L648X, T644X, F627X, S617X, A601X, S575X, LAΔ26, LAΔ26-L648X, LAΔ26-L644X, LAΔ26-L627X, and LC. Terminal amino acids are numbered, and their accumulation dynamics is summarized on the right (+, accumulated at the rupture site; -, not accumulated). **(B** and **C)** The mEmerald-fused LA-FL, PG, PG-CSM, L648X, T644X, F627X, S617X, A601X, S575X and LC were ectopically expressed under the *Lmna* promoter in LA/C-KO cells. **(B)** Representative confocal images of the indicated nuclei. Bar: 5 μm. **(C)** The NP/NL ratios were measured based on mEmerald signals (n = 15-20 cells from two independent experiments; ‡, P<0.001 compared to others except PG-CSM and *, P < 0.05 by a Games–Howell test). **(D** and **E)** Time-lapse images of mEmerald-fused LA-FL, LA mutants, PG, PG deletion mutant and LC were acquired with 10 s intervals for 3 min after the induction of NE rupture by laser-microirradiation, and the relative intensities at the rupture sites are plotted in the graphs. The fluorescence intensities at the rupture sites were measured and normalized to the initial intensities (means ± SEM; n =20 cells from two independent experiments; ***, P < 0.001 from LA-FL by a mixed effect model). **(D)** mEmerald-fused LA-FL, PG, PG-CSM, L648X, T644X, F627X, S617X, A601X, S575X and LC. **(E)** mEmerald-fused LA-FL, LAΔ26, LAΔ26-L648X, LAΔ26-L644X and LAΔ26-L627X. LA-FL and LC are reproduction of D.

## Discussion

A-type lamins are localized to sites of NE rupture for the repair (23, 24, 30). Our previous study shows that endogenous LC but not LA rapidly accumulates the rupture sites within 3 min after the induction of NE rupture because LC is more abundant in the nucleoplasm than LA (30, 39, 40). LA can rapidly accumulate only when LA is overexpressed at a high level to increase the nucleoplasmic pool (30, 37, 38). LC accumulation requires both the immunoglobulin-like fold (Ig-fold) domain that binds to BAF and a nuclear localization signal (NLS), and is not attributable to the LC-specific amino acids (30). In this study, we analyze the LA-specific tail region to determine a molecular mechanism for retaining its small pool in the nucleoplasm and decelerating localization kinetics at the rupture sites, and Fig. 6 illustrates our current findings following our previous study (30). The proteolytic removal of 15 amino acids containing the farnesylated C-terminal end of pre-LA by Zmpste24 increases the nucleoplasmic pool and accelerates the localization kinetics at the rupture sites within the long-time window (60-70 min) after the induction of NE rupture. In HGPS, pre-PG lacks 50 amino acids containing the second proteolytic cleavage site from the C terminus of pre-LA, leading to the permanent association with the INM, which significantly decreases the nucleoplasmic pool and decelerates the localization kinetics at the rupture sites. In contrast, FTI treatment pronouncedly increases the nucleoplasmic pools of LA and PG and promotes their localization to the rupture sites. Therefore, the processes for increasing the nucleoplasmic pool is likely to be rate-limiting steps for the localization kinetics within this time window. Additionally, LACS1 cooperates with LACS2 to inhibit the rapid accumulation at the rupture sites within the short-time window (3 min).

**Figure 6.**
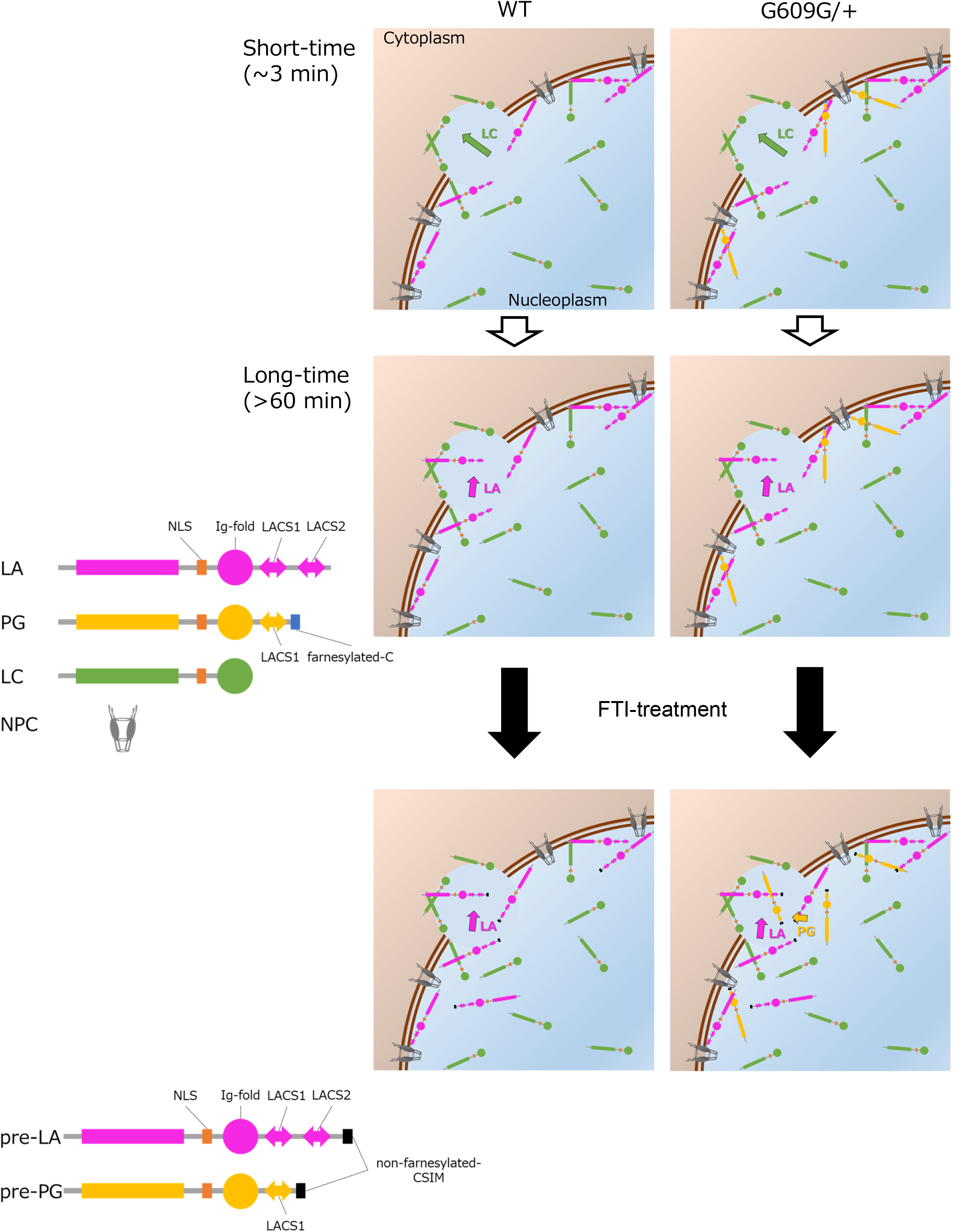
Schematic diagram of the difference in localization to the rupture sites between LA, LC and PG. LA (magenta) is slowly localized to the rupture sites compared to LC (green) because it contains two LACSs in the specific tail region (top and middle). PG (yellow) is even slower in localizing to the rupture sites than LA because of the permanent association with the INM. PG expression also slows the recruitment of LA and LC to the rupture sites. When the farnesylation of PG is inhibited by FTI treatment, the localization kinetics becomes similar to that of LA (bottom).

We use immortalized WT and G609G/+ MEFs to analyze the NP/NL ratios of LA, pre-LA and PG, and their localization kinetics at the rupture sites. Unlike HGPS-patient cells and mouse SMCs expressing PG (32, 34, 58), our G609G/+ cells do not exhibit the formation of misshapen nuclei and spontaneous NE rupture, which could be due to the immortalization process with SV40 large T antigen. In oncogene-induced senescence of normal human fibroblasts, DNA protrusion from the nucleus, micronuclei formation and the leakage of nuclear contents are coupled to the loss of LB1 (59-61). The LB1 expression level could also decrease when HGPS-patient cells become prematurely senescent (62) and PG expression levels increase with age in aortas from G609G/+ mice whereas LB1 levels decrease (34). Therefore, spontaneous NE rupture might be induced by the HGPS mutation because of the induction of premature senescence and rapid cellular aging.

Instead of spontaneous NE rupture, we find that upon the induction of forced NE rupture by AFM indentation, the nuclear reentry of NLS-sfCherry is significantly slow in G609G/+ cells compared to WT cells, strongly suggesting that a repair process of forced NE rupture is impaired by PG expression. The LA expression level in G609G/+ cells is reduced compared to that in WT cells due to the efficient alternative splicing from the mutant allele of the *Lmna* gene (41, 63), which could explain its smaller pool in the nucleoplasm compared to WT cells to decelerate the localization kinetics at the rupture sites. On the other hand, the LC expression level could be increased in G609G/+ cells compared to WT cells for some reason (40, 41). However, the diffusion of the fast component is significantly slow in the nucleoplasm of G609G/+ nuclei compared to WT nuclei, as our results indicate. This could be explained if there is a possibility that LC is increased in the NL and decreased in the nucleoplasm because of the strong association with the NL by PG expression. To support this idea, LC has a significantly slow turnover in the NL of HGPS-patient nuclei compared to normal nuclei (53, 64). These results suggest that the alteration in expression and distribution of LA/C by PG expression impair the repair of NE rupture.

We find that when the farnesylation at the CaaX motif of pre-LA and pre-PG is inhibited by FTI treatment, they are increased and become diffusible in the nucleoplasm to accelerate the localization kinetics to the rupture sites, indicating that the farnesylation process limits the localization to the rupture sites regardless of the second cleavage of pre-LA. Furthermore, Zmpste24 KD blocks the second cleavage, leading to the complete inhibition of pre-LA localization to the rupture sites, and the effects of Zmpste24 KD on the NP/NL ratios and the localization kinetics are rescued by FTI treatment. These results strongly suggest that pre-LA is mostly farnesylated in Zmpste24 KD cells and that the nucleoplasmic pool is little or none to localize to the rupture sites within 60 min.

Apart from the farnesylation at the CaaX motif and the inhibition of the second cleavage of pre-LA, we find that LACS1 and LACS2 in the tail region are also involved in slowing the localization to the rupture sites. Because the truncation of LACS1 significantly increases the NP/NL ratios, LA appears to be retained at the NL through LACS1 compared to LC, resulting in the slow localization. It is possible that LACS1 contributes to the formation of lamin filaments in the NL and/or interacting with other NE components including INM proteins. On the other hand, LACS2 could assist the LACS1 function for inhibiting LA localization to the rupture sites without promoting the retention of LA at the NL. Because the truncation of the residues 599-625 between LACS1 and LACS2 in human increases the NP/NL ratios, a segment within these amino acids is likely to retain its small pool in the nucleoplasm. Whereas the residues 411-553 including the NLS and the Ig-fold domain interacts with dsDNAs (67), the residues 506-638 binds histone H3 (68). This chromatin-binding domain contains the LACSs, suggesting that LA is partially tethered to chromatin through LACSs to slow the localization to the rupture sites.

In HGPS, the permanent association of PG with the INM by the farnesylation at the CaaX motif significantly reduces the nucleoplasmic pool, leading to the slow localization to the rupture sites. Similarly, this mechanism could be also critical for their accesses to active enhancers in the nuclear interior to affect the gene expression in HGPS patient cells (69). Other studies show that the PG association with the INM can lead to the increased NE instability (32, 34) but we do not observe spontaneous NE rupture in our G609G/+ cells. Nevertheless, our findings of the slow localization kinetics of LA, LC and PG at the rupture sites implicate pathological significance of the defective repair of the NL. The therapeutic effect of FTI treatment appears to rescue from the repair defect by targeting the farnesylation of PG. Therefore, the induction of spontaneous NE rupture and the rescuing with FTI treatment need validating using the mouse model of HGPS in future studies.

## Materials and methods

### Plasmid construction

Two different NLSs derived from SV40 large T antigen (NLS^SV40^; PKKKRKV) and c-Myc (NLS^Myc^; PAAKRVKLD) from pCDH-NLS^SV40^-sfCherry-NLS^Myc^-Blast (30) were ligated to pCDH-CMV-MCS-EF1-Hygro (plasmid #CD515B-1, System Biosciences) using the Ligation-Convenience Kit (Nippon Gene) and designated as pCDH-NLS^SV40^- sfCherry-NLS^Myc^-Hygro. The annealed oligonucleotides to express shRNAs (Zmpste24, TRCN0000366641; BAF#1, TRCN0000124958; BAF#2, TRCN0000124955; Table S2, Broad Institute) were ligated into pLKO.3-blast (30) using the Ligation-Convenience Kit. The pX459-sfCherry CRISPR-Cas9 vector were generated from pSpCas9(BB)-2A-Puro (pX459) V2.0 (Addgene plasmid # 62988; http://n2t.net/addgene:62988; RRID:Addgene_62988; a gift from Feng Zhang). The annealed oligonucleotides to express single-guide RNA (sgRNA) against the *Rosa26* target sequence (70) was ligated into pX459-sfCherry using the Ligation-Convenience Kit and designated as pX459-sfCherry-sgRosa26-1.

We cloned a 1.6 kb genomic region upstream from start codon of mouse *Lmna* as the endogenous promoter (-1407 to +249 bp from transcription start site) using PrimeSTAR HS DNA Polymerase (Takara). Primers used in this study are listed in Table S2. Mouse genomic DNA was extracted from cells by Proteinase K (Fungal; Invitrogen) digestion according to the standard protocol and used for a PCR template. The CMV promoter of mEmerald-C1 (Addgene plasmid # 53975; http://n2t.net/addgene:53975; RRID:Addgene_53975; a gift from Michael Davidson) was replaced by the *Lmna* promoter using the In-Fusion HD Cloning Kit (Clontech) and designated as pLmna-mEmerald-C1. The mouse pre-LA cDNA was amplified from pPyCAG-LA-IP (71) using the KOD One PCR Master Mix -Blue-(Toyobo), cloned into pLmna-mEmerald-C1 using the In-Fusion, and designated as pLmna-Em-LA. The mouse PG cDNA was generated by PrimeSTAR Max DNA Polymerase (Takara) to skip 150 bp, closed by In-Fusion HD, and designated as pLmna-Em-PG. The amino acid truncation mutants were generated by PCR mutagenesis using PrimeSTAR Max or PrimeSTAR HS DNA Polymerase. The LACS1 deletion mutant (Δ26) was generated using PrimeSTAR Max to skip 78 bp, and closed by In-Fusion HD. The pLmna-Em-LA and pLmna-Em-PG expression cassettes were amplified by the KOD One, fused with bGHpA amplified from LSL-Cas9-Rosa26TV (Addgene plasmid # 61408; http://n2t.net/addgene:61408; RRID:Addgene_61408; a gift from Feng Zhang) and *Pac* I/*Sma* I digested LSL-Cas9-Rosa26TV by In-Fusion HD, and designated as pLmna-Em-LA-R26Neo, pLmna-Em-PG-R26Neo and pLmna-Em-PG-CSM-R26Neo. The mouse LC cDNA was also generated by KOD One PCR and In-Fusion HD and designated as pLmna-Em-LC-R26Neo.

### Cell culture

MEFs isolated from WT and heterozygous c.1827C>T mutant (*Lmna*^G609G/+^, G609G/+) embryos at embryonic day 13.5 were generously gifted by Carlos López-Otín (Universidad de Oviedo) (41, 42). Primary MEFs were cultured in modified DMEM (high glucose; Nacalai Tesque) containing 10% FBS (qualified; Thermo Fisher), 4 mM L-glutamine, 100 U/mL penicillin and 100 μg/mL streptomycin (Sigma-Aldrich), and 20 mM 2-[4-(2-Hydroxyethyl)-1-piperazinyl] ethanesulfonic acid (HEPES; Nacalai Tesque) at 37°C in a humidified multi-gas incubator with 3% O_2_/5% CO_2_ (MCO-5M, Sanyo/Panasonic Healthcare). Cells were immortalized by transduction with a retrovirus expressing SV40 large T antigen from pBABE-puro largeTcDNA (Addgene plasmid # 14088; http://n2t.net/addgene:14088; RRID:Addgene_14088; a gift from William Hahn) as previously described (13). After selection with 2 μg/ml puromycin (InvivoGen) for 4 d, immortalized WT and G609G/+ cells were cultured in growth medium without HEPES at 37°C in a humidified chamber with 5% CO_2_.

### Lentiviral transduction

For lentivirus-mediated stable introduction of NLS^SV40^-sfCherry-NLS^Myc^, we followed the methods that was previously described (30). Briefly, pVSV-G (PT3343-5, Clontech) and psPAX2 (Addgene plasmid #12260; http://n2t.net/addgene:12260; RRID:Addgene_12260; a gift from Didier Trono), together with a pCDH vector (pCDH-NLS^SV40^-sfCherry-NLS^Myc^-Blast, pCDH-NLS^SV40^-sfCherry-NLS^Myc^-Hygro or pCDH-sfGFP-DARPin-LA6-Hygro) in a 1:3:4 weight ratio of each plasmid was transfected into ∼80% confluent 293T cells (CRL-3216, ATCC) using Lipofectamine 3000 following the manufacturer’s instructions for lentivirus production. 24 h after the transfection, the medium was replaced with fresh medium, and the virus-containing culture supernatant was harvested at 48 h after transfection and filtrated with a 0.45-μm syringe filter. For virus infection, MEFs were incubated in the culture supernatants with 4 µg/mL polybrene (Nacalai Tesque) for 24 h. Infected cells were selected by incubation in medium containing 3 µg/mL blasticidin S or 200 µg/mL hygromycin B Gold (InvivoGen) for 4 d.

### Live-cell imaging and NE rupture induction by laser microirradiation

Culture medium was replaced with FluoroBrite DMEM (Thermo Fisher) containing 10% FBS, 4 mM L-glutamine, 100U/mL penicillin and 100 μg/mL streptomycin. For laser-microirradiation and image acquisition, cells were set onto a heated stage (37°C; Tokai Hit) with a CO_2_-control system (Tokken) on a confocal microscope (FV1000, Olympus) with a 60× PlanApo N (NA 1.4) oil lens operated by built-in software FLUOVIEW ver. 4.2 (Olympus). All live-cell images were acquired using a main scanner (4% 488-nm laser transmission; 30% 543-nm laser transmission; 2 μs/pixel; 512×512 pixels; pinhole 100 μm; 6× zoom; ∼10 s/frame). After the first image was acquired, a 2 μm diameter spot was laser-microirradiated using a second scanner at 100% power of 405-nm laser transmission (40 μs/pixel; approximately 780 μW) for 10 s whereas the images were acquired with another scanner.

Fiji software (ImageJ) ver. 1.53v (National Institute of Health) was used for measuring max intensity at the sites of NE rupture after Gaussian filtering with σ = 2.0 over time. Fluorescence intensity was normalized by each of the initial intensity in the laser-microirradiated region.

CellProfiler ver. 3.1.9 (Broad Institute) with the NP_NL_intensity_live pipeline (uploaded on GitHub; http://github.com/Kimura-Lab/Kono-et-al.-2023) (30) were used for measuring mean intensity of mEmerald-LA mutants and mEmerald-LC in the NP and the NL. Briefly, after the nuclear region was recognized by the nuclear localization of NLS-sfCherry, 3 pixels (207 nm) and more than 10 pixels (690 nm) inside from the rim of nuclear region were regarded as the NL and the NP, respectively, and then the mean intensity of each section was measured. All images were processed in Photoshop ver. 24 (Adobe) for cropping and brightness/contrast adjustment.

### Indirect immunofluorescence and microscopy

Primary antibodies used for immunofluorescence were mouse monoclonal anti-LA (1:250; 4A58, sc-71481, Santa Cruz), rat monoclonal anti-pre-LA (1:1,000; 7G11; MABT345, EMD Millipore; (49)), mouse monoclonal anti-mature-LA (1:8,000; 4A4-A4, 14-9848-82, Invitrogen; (72)), rabbit polyclonal anti-progerin (1:1000; generated by the Nourshargh laboratory using a peptide immunogen of murine progerin and standard immunization procedures; (63, 73, 74)), mouse monoclonal anti-LB1 (1:1,000; B-10; sc-374015, Santa Cruz), rabbit monoclonal anti-cGAS (1:250; D3O8O; 21659, Cell Signaling Technology) and rabbit monoclonal anti-BANF1/BAF (1:200; EPR7668; ab129184, Abcam). The secondary antibodies used were Alexa Fluor 488-donkey anti-mouse immunoglobulin G (IgG), Alexa Fluor 488-donkey anti-rabbit IgG (1:2,000; A21202 and 1:2,000; A21206, respectively, Thermo Fisher), Cy5-donkey anti-mouse IgG and Cy5-goat anti-rat IgG (1:2,000; 715-175-151 and 1:2,000; 712-175-153, respectively, Jackson ImmunoResearch).

Cells were grown on 35-mm Glass Based Dishes which have a cover glass with a grid pattern (IWAKI) and fixed after laser-microirradiation with 4% PFA (Electron Microscopy Sciences) containing 0.1% Triton X-100 (Nacalai Tesque) for 15 min, followed by permeabilization using 0.1% Triton X-100 in PBS for 10 min and then blocking with Blocking One-P (Nacalai Tesque) for 30 min. Cells were incubated with primary antibodies overnight at 4°C, washed with 10% Blocking One-P in PBS, and incubated with secondary antibodies for 1 h at RT. DNA was stained with Hoechst 33342 (0.5 μg/mL; Thermo Fisher). For Fig. S1 A, cells were fixed with 4% PFA for 15 min, followed by permeabilization using 0.1% Triton X-100 in PBS for 10 min and then blocking with Blocking One-P for 30 min. After incubation with primary antibodies, the RFP-Booster Alexa Fluor 568 (alpaca monoclonal anti-RFP Nanobody; 1:1000; rb2AF568, ChromoTek) was used with second antibodies to amplify the signal of NLS-sfCherry.

Confocal fluorescence images were obtained using a confocal microscope FV1000 with a 60× PlanApo N (NA 1.4) oil lens (3.0% 405-nm laser transmission; 3.1-11.1% 488-nm laser transmission; 5.0% 543-nm laser transmission; 1.0-4.0% 633-nm laser transmission; 2 μs/pixel; 512×512 pixels; pinhole 100 μm). CellProfiler with the NP_NL_intensity_fixed pipeline (uploaded on GitHub; http://github.com/Kimura-Lab/Kono-et-al.-2023) were used for measuring mean intensity of lamins in the NP and the NL. The nuclear region was recognized by staining DNA with Hoechst 33342.

### Induction of mechanical NE rupture by atomic force microscopy (AFM)

Culture medium was replaced with Leibovitz’s L-15 medium (no phenol red; Thermo Fisher) containing 10% FBS, 2 mM L-glutamine, 100U/mL penicillin and 100 μg/mL streptomycin. JPK NanoWizard 4 BioAFM (Bruker) incorporated into an inverted fluorescence microscope (Eclipse Ti2, Nikon) with EM-CCD camera (iXon Ultra 888, Andor) and light source (X-Cite XYLIS, Excelitas) was used for single-cell compression. Cantilever (TL-CONT, Nanosensors; spring constant approximately 0.2 N/m) with a 100-μm glass bead (BZ-01, As-One) was used for NE rupture. Cells expressing NLS-sfCherry were set onto a heated stage (37°C) and the first fluorescent image of NLS-sfCherry was obtained before the rupture using a TRITC filter and NIS-Elements BR (Nikon). Next, the bead was approached on the top of the cell surface with 2 nN setpoint and then forced down for 5 μm. After waiting for 10 s, the cantilever was retracted. Within 10 s after the retraction, time-lapse imaging was started with 30 s exposure time every 30 s for 5 min. Fiji software (ImageJ) was used for measuring mean intensities in the region of interest from nucleoplasm and cytoplasm.

### Chemicals

As a selective farnesyltransferase inhibitor (FTI), lonafarnib (SCH66336; Merck) was dissolved in dimethylsulfoxide (DMSO; Nacalai Tesque) and added to the cells with one daily dose of 3.2 μM for 2 days. For controls in all experiments, an equal volume of vehicle (0.1% DMSO) was added.

### Immunoblotting

For immunoblotting, we followed the methods that were previously described (30). Briefly, primary antibodies used for immunoblotting were mouse monoclonal anti-LA/C (1:5,000-20,000; 3A6-4C11, 14-9847-82, eBioscience), rat monoclonal anti-pre-LA (1:1,000; 7G11), rabbit polyclonal anti-progerin (1:1,000; Nourshargh lab), rabbit monoclonal anti-BANF1/BAF (1:500; EPR7668, Abcam), anti-β-Actin (1:2,000; AC-15, sc-69879, Santa Cruz) and anti-GAPDH (1:5,000; 6C5, sc-32233, Santa Cruz). The secondary antibodies used were anti-mouse IgG HRP-linked whole Ab sheep (1:10,000; NA931, Amersham), anti-rabbit IgG HRP-linked F(ab’)_2_ fragment donkey (1:10,000; NA9340, Amersham) and HRP-donkey anti-rat IgG (H+L) (1:10,000; 712-035-150, Jackson ImmunoResearch).

### Rosa26-targeted gene knock-in

The pLmna-Em-preLA-R26Neo, pLmna-Em-PG-R26Neo and pLmna-Em-LC-R26Neo were linearized by the *Kpn* I (Takara) and purified by phenol/chloroform extraction and ethanol precipitation. Cells were transfected with appropriate plasmids using Lipofectamine 3000 (Thermo Fisher) by reverse transfection as previously described (30). Briefly, the DNA-lipid complex (3 μg linearized *Rosa26* targeting vectors : 2 μg pX459-sfCherry-sgRosa26-1 : 7.5 μL Lipofectamine 3000 in 250 μL Opti-MEM; Thermo Fisher) was added to 1.62×10^5^ cells as they were seeding onto a 35 mm dish with 2 mL of growth medium. One day after the transfection, the medium was replaced with fresh medium. Two days after the transfection, cells were selected by incubation in medium containing 400 µg/mL G-418 disulfate aqueous solution (Nacalai Tesque) for 7 d.

### Fluorescence Correlation Spectroscopy (FCS)

FCS measurements were all performed at 25°C with an LSM780 confocal microscope (Carl Zeiss) as previously described (15, 46, 75). FCS setups using the LSM780 microscope consisted of a continuous-wave Ar^+^ laser (25 mW), a water-immersion objective (C-Apochromat, ×40/1.2 NA; Carl Zeiss), and a GaAsP multichannel spectral detector (Quasar; Carl Zeiss). Fluorescent protein of mEmerald was excited with the 488-nm laser line with a minimal laser power (0.5%) to allow an optimal signal-to-noise ratio. Diameter of pinhole was adjusted to 34 μm (1 Airy unit) for the 488-nm laser. Emission fluorescence was detected at 500–550 nm in the green channel. The structure parameter and detection volume were calibrated using Rhodamine-6G (Sigma-Aldrich) solution with diffusion coefficient (280 μm^2^/s; 25°C in water) before experiments with cells. For FCS measurement, three positions were randomly selected inside the nucleoplasm, and molecular diffusion was measured for 10 s five times. The first round of the five repetitions were carried out to photobleach immobile fraction, and then the four repetitions were used for further analysis. FCS data were analyzed with analytical software installed in the ZEN 2012 SP5 FP1 (black) acquisition software (Carl Zeiss) (46, 75). Briefly, all measured fluorescence auto-correlation functions (FAFs) from live cells were globally fitted by the ConfoCor3 software installed on LSM780 system using fitting models considering three-dimensional free diffusion with a Gaussian laser profile, including a triplet component to estimate the diffusion coefficient.

### Generation of the Zmpste24- and BAF-KD MEFs

MEFs expressing shRNAs of anti-Zmpste24 and anti-BAF were generated through lentiviral transduction. KD by shRNA expression was carried out by Lentivirus transduction (see the section on Lentiviral transduction above) and 3 d selection with 3 µg/mL blasticidin S Gold for pLKO.3-blast.

### Statistical analyses

Unless mentioned otherwise, all plots showed mean values ± SEM (error bars). Fisher’s exact test was performed to analyze the categorical data using EZR on R commander ver. 1.61 (programmed by Yoshinobu Kanda) (76). The mixed effect model fit by restricted maximum likelihood (REML) was performed to analyze the interaction between group and time using the *lme4* (77) and the *lmerTest* (78) packages in EZR, where cell’s id was set as a random effect, while group and time as fixed effects. The interaction between group and time (group × time) was also set as a fixed effect. The Welch’s t-test or the Welch’s one-way ANOVA followed by the Games–Howell post-hoc test was performed to single or multiple comparison, respectively. Statistic codes for R ver. 4.2.2 and a modified R commander for EZR 1.61 are uploaded on GitHub (http://github.com/Kimura-Lab/Kono-et-al.-2023). Significance was determined if P < 0.05.

## Supplemental material

Table S1 shows the phenotype characterization in WT and G609G/+ cells by immunofluorescence. Table S2 shows oligonucleotide sequences used in this study. Fig. S1 shows the immunofluorescence of WT and G609G/+ cells for phenotype characterization, and the localization of LA and PG to the rupture sites within 10 min, and at 30-40 and 120-130 min after laser microirradiation. Fig. S2 shows the protein architecture of A-type lamins and DARPin-LA6, and localization kinetics of DARPin-LA6 at the rupture sites in WT and G609G/+ cells. Fig. S3 shows WT and G609G/+ cells expressing A-type lamins under the *Lmna* promoter knocked-in to *Rosa26* locus and FCS analysis for mEmerald-LC. Fig. S4 shows the localization of LA and PG to the rupture sites within 10 min in FTI-treated WT and G609G/+ nuclei. Fig. S5 shows the immunoblotting of FTI-treated WT and G609G/+ cells expressing A-type lamins under the *Lmna* promoter knocked-in to *Rosa26* locus, and the representative image of FCS analysis. Fig. S6 shows immunoblotting, immunofluorescence and the localization of LA and pre-LA to the rupture sites in Zmpste24-KD cells. Fig. S7 shows the differences between LA and LC amino acid sequence, and dynamics of LA mutants at the rupture sites, and the NP/NL ratios of mEmerald-LA mutants by live-cell imaging.

## Acknowledgments

We would like to thank Prof. Carlos López-Otín (Universidad de Oviedo) for providing us with the primary MEFs isolated from WT and *Lmna* c.1827C>T mutant (*Lmna*^G609G/+^ and *Lmna*^G609G/G609G^) embryos. We thank Biomaterials Analysis Division, Open Facility Center (OFC) of Tokyo Institute of Technology for nucleotide sequencing and sharing research equipment, especially Carl Zeiss ElyraS1/Airyscan/LSM780 is shared in MEXT Project Grant Number JPMXS0440200022. We also thank the Cellular Imaging Core Facility at the ConveRgence mEDIcine research cenTer (CREDIT), Asan Medical Center for technical support. The generation and validation of the anti-murine progerin antibody was funded by the Wellcome Trust (098291/Z/12/Z to S. Nourshargh) and the British Heart Foundation (FS/IBSRF/22/25121 to L. Rolas). We also acknowledge the R statistic code generated by Shinsuke Hatano (Ryukoku University). We also thank all members of the Kimura lab at Tokyo Institute of Technology and Tokuko Haraguchi (Osaka University) for advice and discussions. This work was supported by JSPS KAKENHI Grant Numbers JP20KK0158 (to T. Shimi, Y. Kono, K. Miyazawa and H. Kimura), JP20K06617 (to T. Shimi), and JP18H05527 and JP21H04764 (to H. Kimura). This work was also supported by MEXT (World Premier International Research Center Initiative [WPI]). Y. Kono was an academy scholar at the Specialized Academy for Cell Science (Ohsumi-juku) of the Tokyo Tech Organization for Fundamental Research (OFR). The authors declare no competing financial interests.

## Author contributions

T. Shimi conceived the project. Y. Kono designed, performed and interpreted experiments. C.G. Pack performed FCS experiments. T. Ichikawa performed AFM experiments. A. Komatsubara performed image analysis. L. Rolas and S. Nourshargh generated, characterized, and provided the anti-murine progerin antibody. S.A. Adam, K. Miyazawa, O. Medalia and R.D. Goldman provided advice and scientific discussion. Y. Kono, C.G. Pack, T. Ichikawa and T. Shimi drafted the manuscript. S.A. Adam, R.D. Goldman, and H. Kimura edited the manuscript. T. Shimi, S.A. Adam, T. Fukuma, H. Kimura and R.D. Goldman supervised the experiments.

## Abbreviations

BAF: barrier-to-autointegration factor
cGAS: cyclic GMP-AMP synthase
DARPin: designed ankyrin repeat protein
DCM: dilated cardiomyopathy
ESCRT-III: endosomal sorting complex required for transport-III
ER: endoplasmic reticulum
FAF: fluorescence auto-correlation function
FCS: fluorescence correlation spectroscopy
FTI: farnesyltransferase inhibitor
FPLD: familial partial lipodystrophy
HGPS: Hutchinson-Gilford Progeria syndrome
Ig-fold: immunoglobulin-like fold
INM: inner nuclear membrane
KD: knockdown
KO: knockout
LA: lamin A
LACS: LA-characteristic sequence
LB1: lamin B1
LB2: lamin B2
LC: lamin C
LINC: linker of nucleoskeleton and cytoskeleton
MEF: mouse embryonic fibroblast
MD: muscular dystrophy
NE: nuclear envelope
NL: nuclear lamina
NP: nucleoplasmic pool
NPC: nuclear pore complex
ONM: outer nuclear membrane
PG: progerin
REML: restricted maximum likelihood
sfGFP: superfolder GFP
sfCherry: superfolder Cherry
shRNA: short hairpin RNA
STING: stimulator of interferon genes
WT: wild type

---

**Table S1.** Phenotype characterization of WT and G609G/+ MEFs. NLS-sfCherry expressing WT and G609G/+ cells were stained with Hoechst 33342 for DNA and RFP-Booster Alexa Fluor 568 to enhance the signal of NLS-sfCherry to detect spontaneous NE rupture. Immunostaining with α-LB1 and α-cGAS antibodies were also used as NE rupture markers. See Fig. S1 A for the confocal images. P values, Fisher’s exact tests between WT and G609G/+.

|  | WT<br>(n = 118 cells,<br>129 nuclei) | G609G/+<br>(n = 116 cells,<br>131 nuclei) | P value |
| --- | --- | --- | --- |
| DNA |  |  | 0.465 |
| normal | 122 (94.6%) | 120 (91.6%) |  |
| misshapen | 7 (5.4%) | 11 (8.4%) |  |
| NLS-sfCherry |  |  | 0.102 |
| nuclear-localized | 110 (93.2%) | 114 (98.3%) |  |
| ruptured | 8 (6.8%) | 2 (1.7%) |  |
| α-LB1 |  |  | 0.212 |
| normal distributed | 125 (96.9%) | 130 (99.2%) |  |
| mis-localized | 4 (3.1%) | 1 (0.8%) |  |
| α-cGAS |  |  | 1 |
| not accumulated | 129 (100%) | 131 (100%) |  |
| accumulated | 0 (0%) | 0 (0%) |  |

**Table S2.**
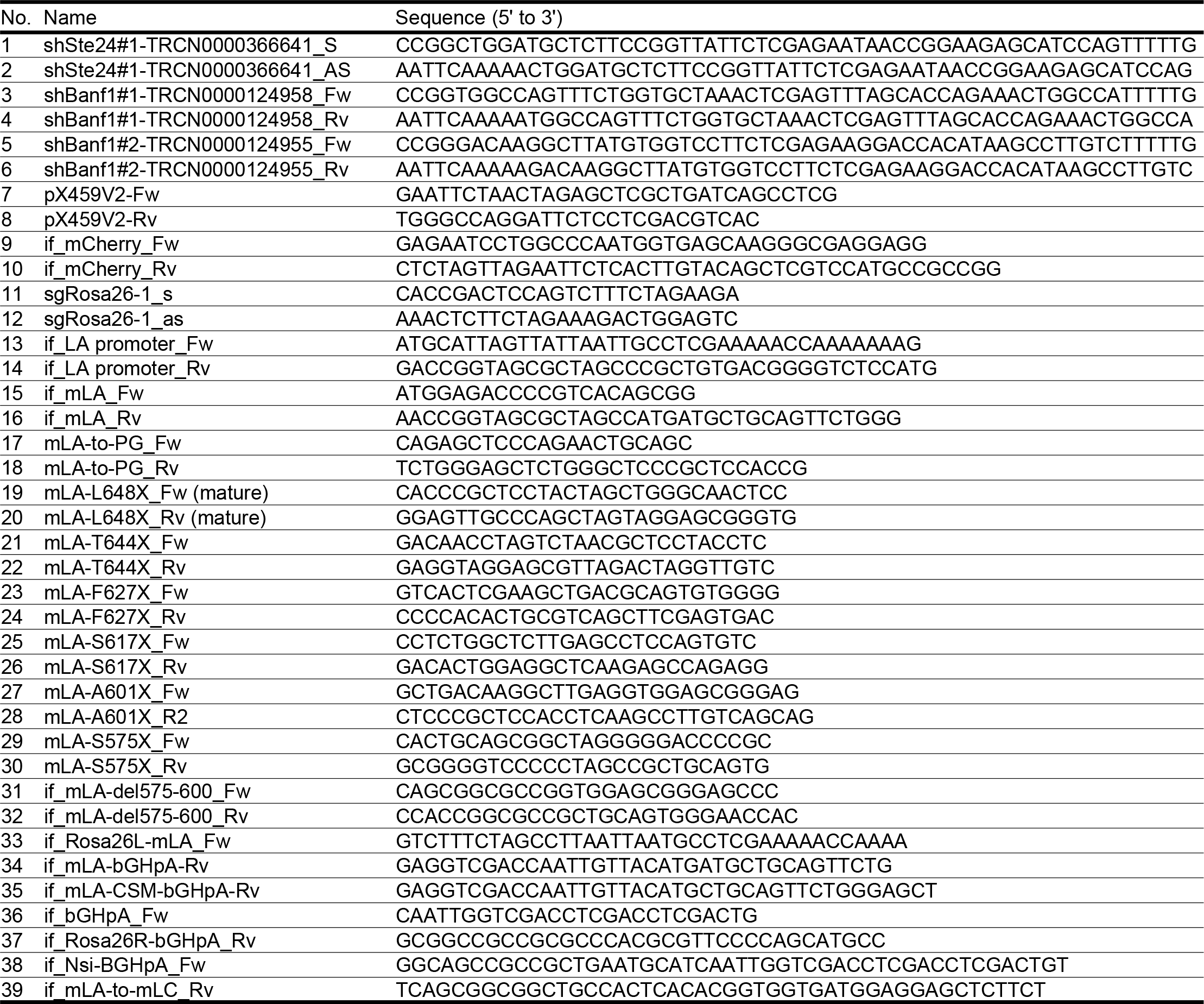
Oligonucleotides used in this study. See Materials and methods. Oligonucleotides were synthesized by Sigma-Aldrich or Integrated DNA Technologies (IDT).

**Figure S1.**
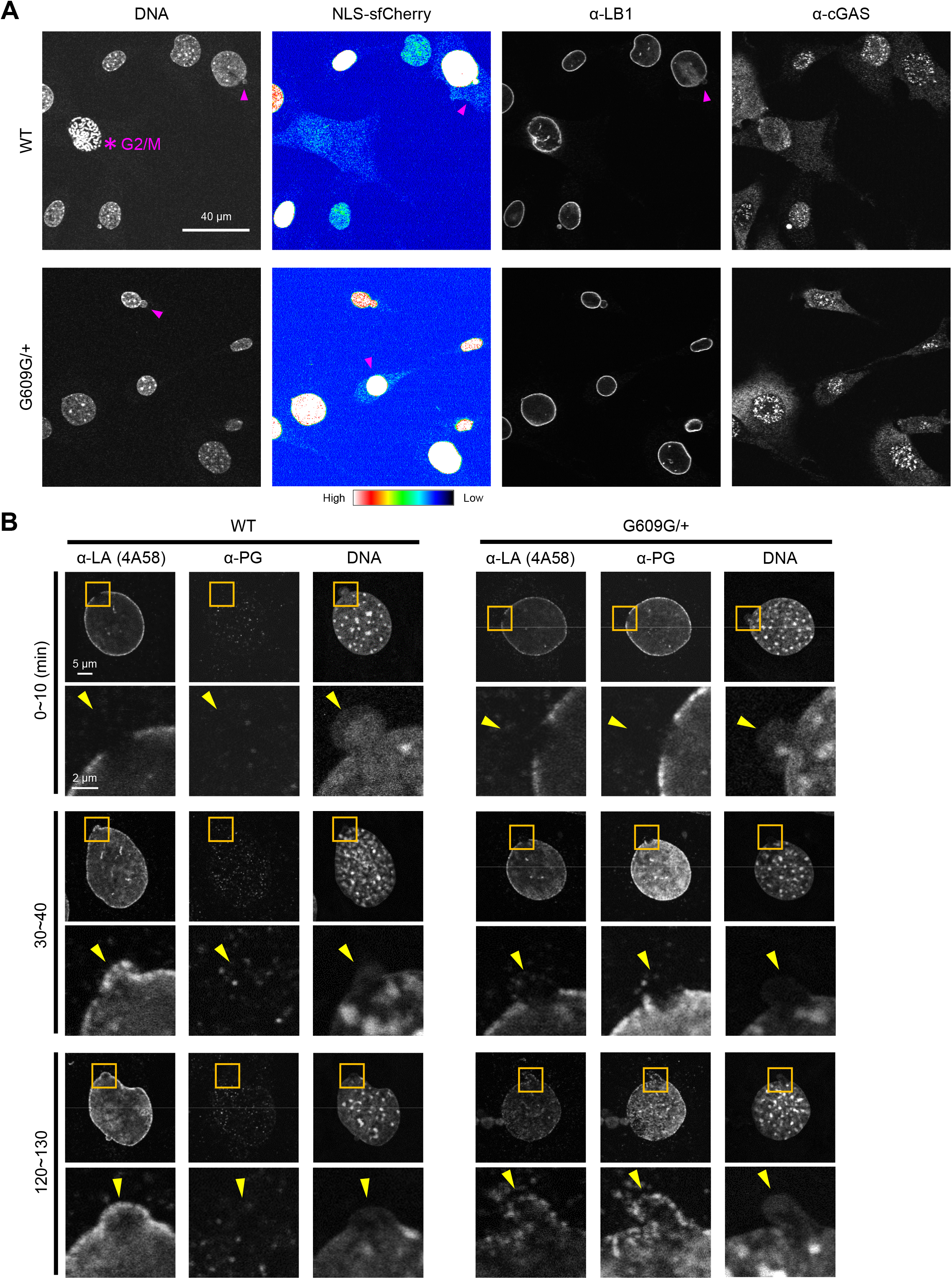
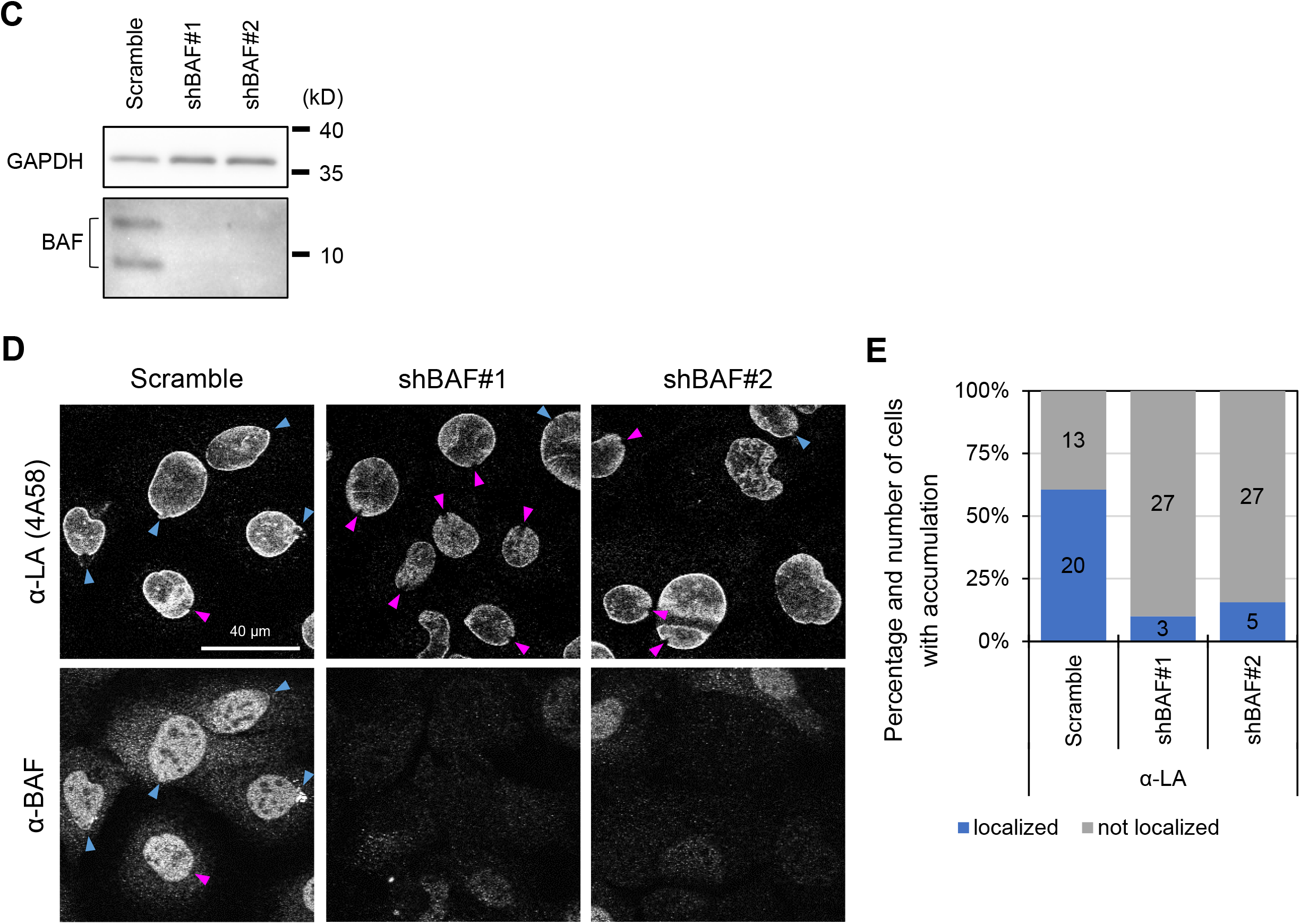
Phenotype characterization, the localization kinetics of LA and PG at the rupture sites in WT and G609G/+ MEFs, and the localization kinetics of LA at the rupture sites in BAF-KD WT MEFs. **(A)** The phenotype of heterozygous knock-in G609G/+ expressing NLS-sfCherry was characterized by immunofluorescence. The cells were immunostained with anti-LB1 and anti-cGAS, and the DNA was stained with Hoechst 33342. The fluorescence intensities of NLS-sfCherry are shown with rainbow color. Asterisk indicates a cell with NE breakdown due to the G2-M transition and were excluded from the analysis. Misshapen, NE-ruptured, LB1-mis-localized and cGAS-accumulated nuclei are indicated with magenta arrowheads. **(B)** Representative confocal images of the nuclei. Magnified views of the indicated areas by orange boxes are shown (bottom of each row). The cells were fixed within 0-10 min (top), 30-40 min (middle) and 120-130 min (bottom) after the induction of NE rupture by laser microirradiation, followed by fixation with 4% PFA/0.1% Triton-X 100. The fixed cells were immunostained with anti-LA (4A58) and anti-PG. The rupture sites are indicated with yellow arrowheads (bottom of each row). Bars: 5 μm (top of each row) and 2 μm (bottom of each row). **(C)** Whole cell lysates from WT cells expressing shRNAs for the control (Scramble) or two shRNAs (shBAF #1 and #2) were probed with anti-BANF1/BAF and anti-GAPDH (as loading control) for immunostaining. Positions of the size standards are shown on the right. **(D** and **E)** 15 to 19 of the indicated cells were laser-microirradiated within 10 min and incubated for 60 min, followed by fixation with 4% PFA/0.1% Triton-X 100. The fixed cells were immunostained with the LA and BAF antibodies. **(D)** Representative confocal images of nuclei with (blue arrowheads) or without (magenta arrowheads) the localization of LA to the rupture sites. Bar: 40 μm. **(E)** Percentiles of cells with (blue) and without (gray) the localization of LA. The numbers of analyzed cells from two independent experiments are indicated in the bar charts.

**Figure S2.**
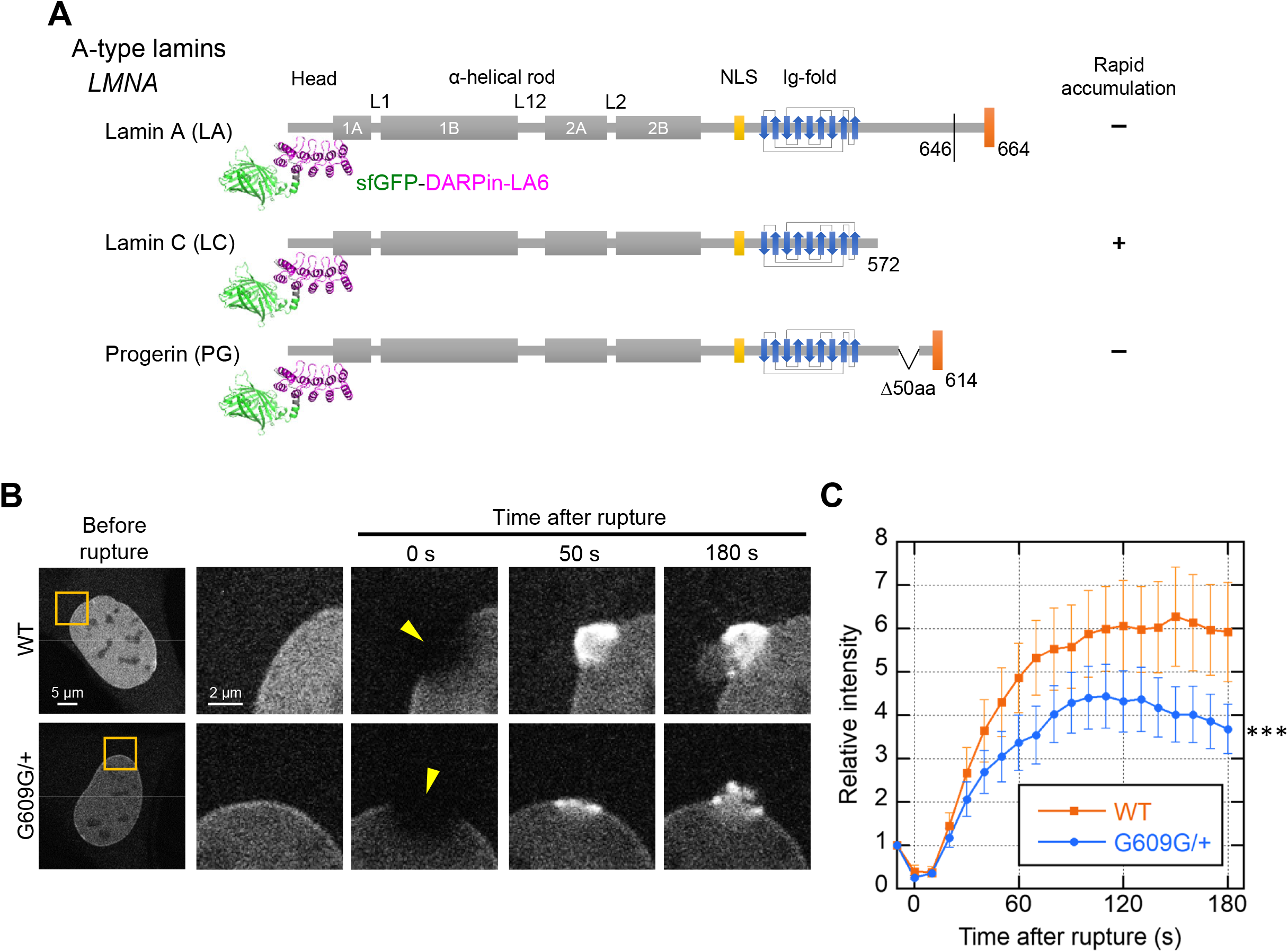
Accumulation kinetics of LC at the rupture sites in WT and G609G/+ MEFs. **(A)** Protein architecture of LA, LC and PG with sfGFP-DARPin-LA6. The summary of their accumulation at the rupture sites is indicated on the right (+, accumulated at the rupture site; -, not accumulated). The structural image of sfGFP-DARPin-LA6 was predicted by AlphaFold2 (79) and edited by the PyMOL (Schrödinger). **(B** and **C)** Time-lapse images of sfGFP-DARPin-LA6 expressed in WT and G609G/+ cells were acquired with 10 s intervals for 3 min after the induction of NE rupture by laser-microirradiation, and the relative intensities are plotted in the graph. **(B)** The dynamics of the sfGFP-DARPin-LA6 in response to NE rupture in the cells. Magnified views of the areas indicated by orange boxes are shown (the second to fifth columns). A 2-μm diameter spot at the NE was laser-microirradiated to induce NE rupture (yellow arrowheads). Bars: 5 μm (the first column) and 2 μm (the second to fifth columns). **(C)** The fluorescence intensities at the rupture sites were measured and normalized to the initial intensities (means ± SEM; n = 20 cells from two independent experiments; ***, P < 0.001 from WT by a mixed effect model).

**Figure S3.**
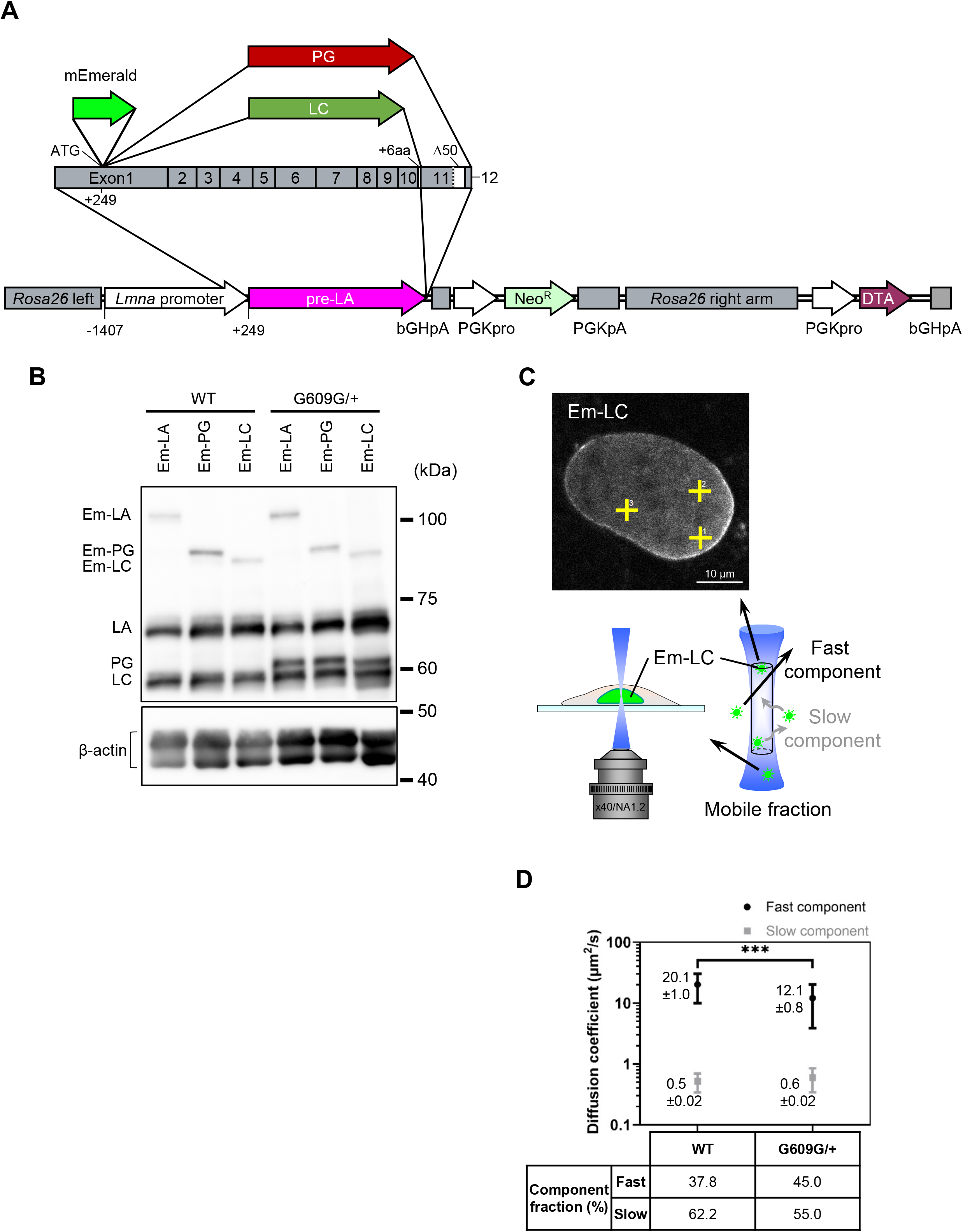
Establishment of knock-in cells expressing mEmerald-fused LA, PG and LC under the *Lmna* promoter for FCS measurements. The DNA sequences of mEmerald-LC, mEmerald-LA and mEmerald-PG with the endogenous *Lmna* promotor were knocked-in to the *Rosa26* locus in WT and G609G/+ cells. **(A)** The schematic diagram of linearized pLmna-Em-LA-R26Neo, pLmna-Em-PG-R26Neo and pLmna-Em-LC-R26Neo knock-in vectors. **(B)** Whole cell lysates from the knocked-in cells with pLmna-Em-LA-R26Neo, pLmna-Em-PG-R26Neo and pLmna-Em-LC-R26Neo were probed with anti-LA/C (3A6-4C11) and anti-β-actin (as loading control) for immunostaining. Positions of the size standards are shown on the right. **(C** and **D)** The diffusion fractions and coefficients of mEmerald-LC (Em-LC) in WT and G609G/+ nuclei were measured by FCS. **(C)** A representative confocal image of a WT nucleus with mEmerald-LC before FCS measurements (top). The yellow crosses indicate the points measured by FCS. Bar: 10 µm. Em-LC molecules move in or out of the confocal volume (white-out cylinder region in blue) at different speeds, as shown in the diagram (bottom). **(D)** Diffusion coefficient of fast component (plotted in black) and slow component (plotted in gray) for Em-LC in the cells. Mean ± SEM are indicated on the left to plots; n = 10 cells from two independent experiments; ***, P < 0.001 by a Welch’s t-test. Percentiles of these components are indicated at the bottom of the graph.

**Figure S4.**
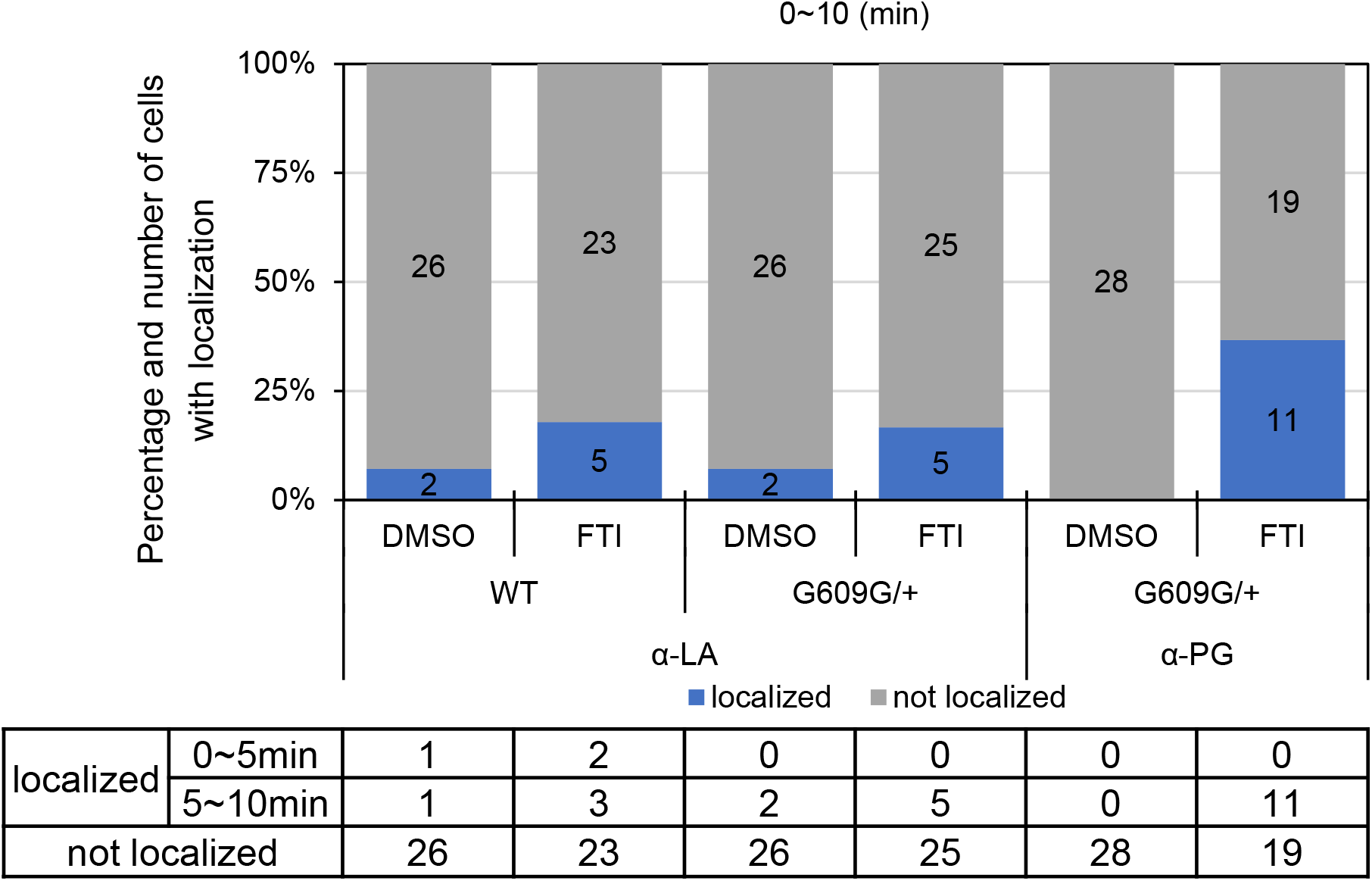
Short-term localization of LA and PG to the rupture sites in DMSO or FTI-treated WT and G609G/+ nuclei. 14 to 15 of nuclei in the cells were laser-microirradiated within 10 min, followed by fixation with 4% PFA/0.1% Triton-X 100. The fixed cells were immunostained with a combination of the LA and PG antibodies. Percentiles of the cells with (blue) and without (gray) the localization of LA and PG at the rupture sites. The numbers of analyzed cells from two independent experiments are indicated in the bar charts. The number of the cells in which LA and PG were localized to the rupture sites are indicated at the bottom of the graph.

**Figure S5.**
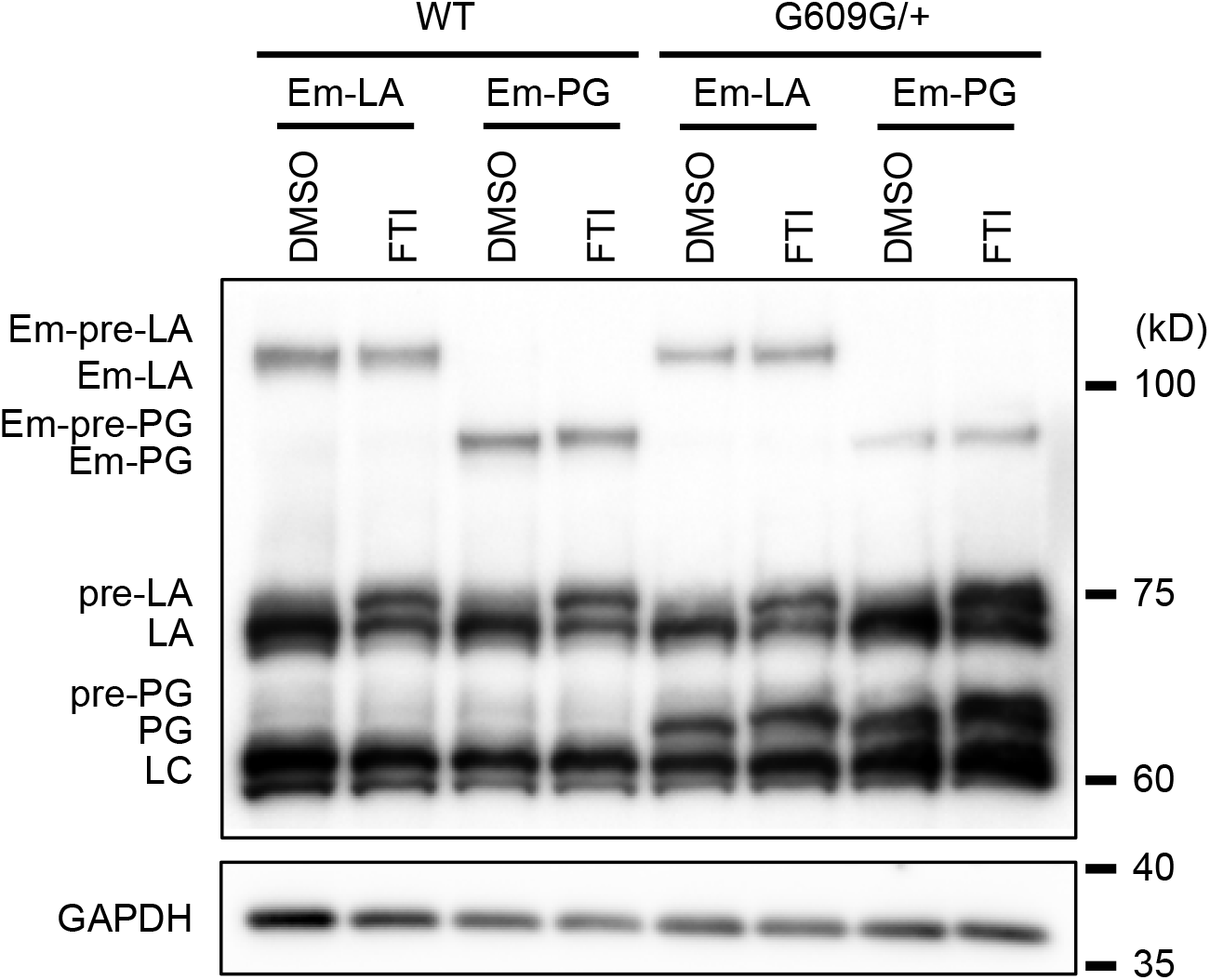
The immunoblotting of FTI-treated WT and G609G/+ knocked-in MEFs. Whole cell lysates from the knocked-in cells with pLmna-Em-LA-R26Neo (Em-LA) and pLmna-Em-PG-R26Neo (Em-PG) treated with DMSO or FTI were probed with anti-LA/C (3A6-4C11) and anti-GAPDH (as loading control) for immunostaining. Positions of the size standards are shown on the right.

**Figure S6.**
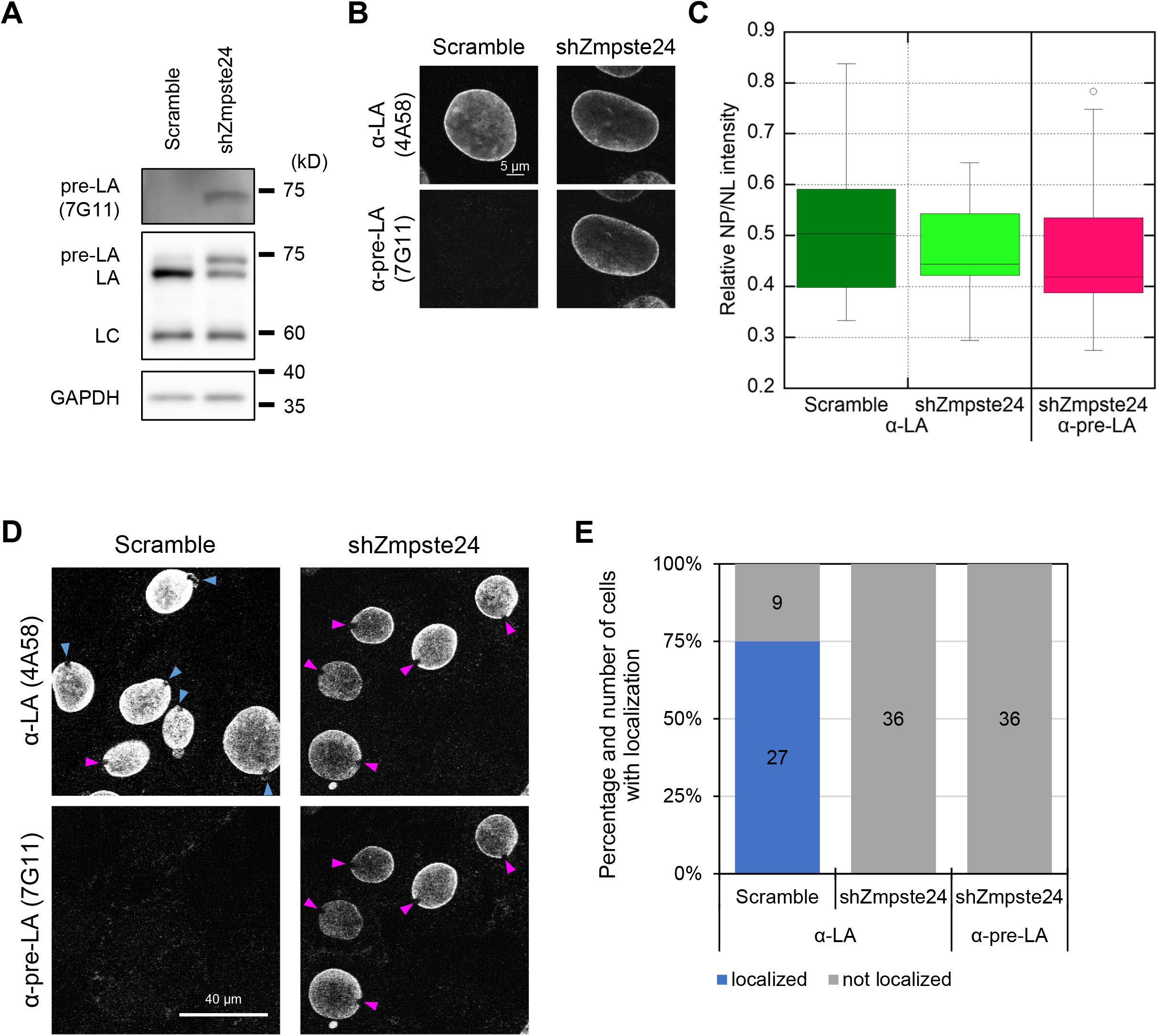
The localization kinetics at the rupture sites in the control and Zmpste24-KD WT MEFs. **(A)** Whole cell lysates from WT cells expressing Scramble for the control or shZmpste24 for Zmpste24-KD were probed with anti-pre-LA (7G11), anti-LA/C (3A6-4C11) and anti-GAPDH (as loading control) for immunobloting. Positions of the size standards are shown on the right. **(B** and **C)** The cells were immunostained with anti-LA (4A58) and the pre-LA (7G11) antibody. **(B**) Representative confocal images of the nuclei. Bar: 5 μm. **(C)** NP/NL ratios of LA and pre-LA were measured based on immunofluorescence (n = 20 cells from two independent experiments). **(D** and **E)** 18 of nuclei in the cells were laser-microirradiated within 10 min and incubated for 60 min, followed by fixation with 4% PFA/0.1% Triton-X 100. The fixed cells were immunostained with the LA and pre-LA antibodies. **(D)** Representative confocal images of the nuclei with (blue arrowheads) or without (magenta arrowheads) the localization of LA and pre-LA to the rupture sites. Bar: 40 μm. **(E)** Percentiles of the cells with (blue) and without (gray) the localization of LA and pre-LA. The numbers of analyzed cells from two independent experiments are indicated in the bar charts.

**Figure S7.**
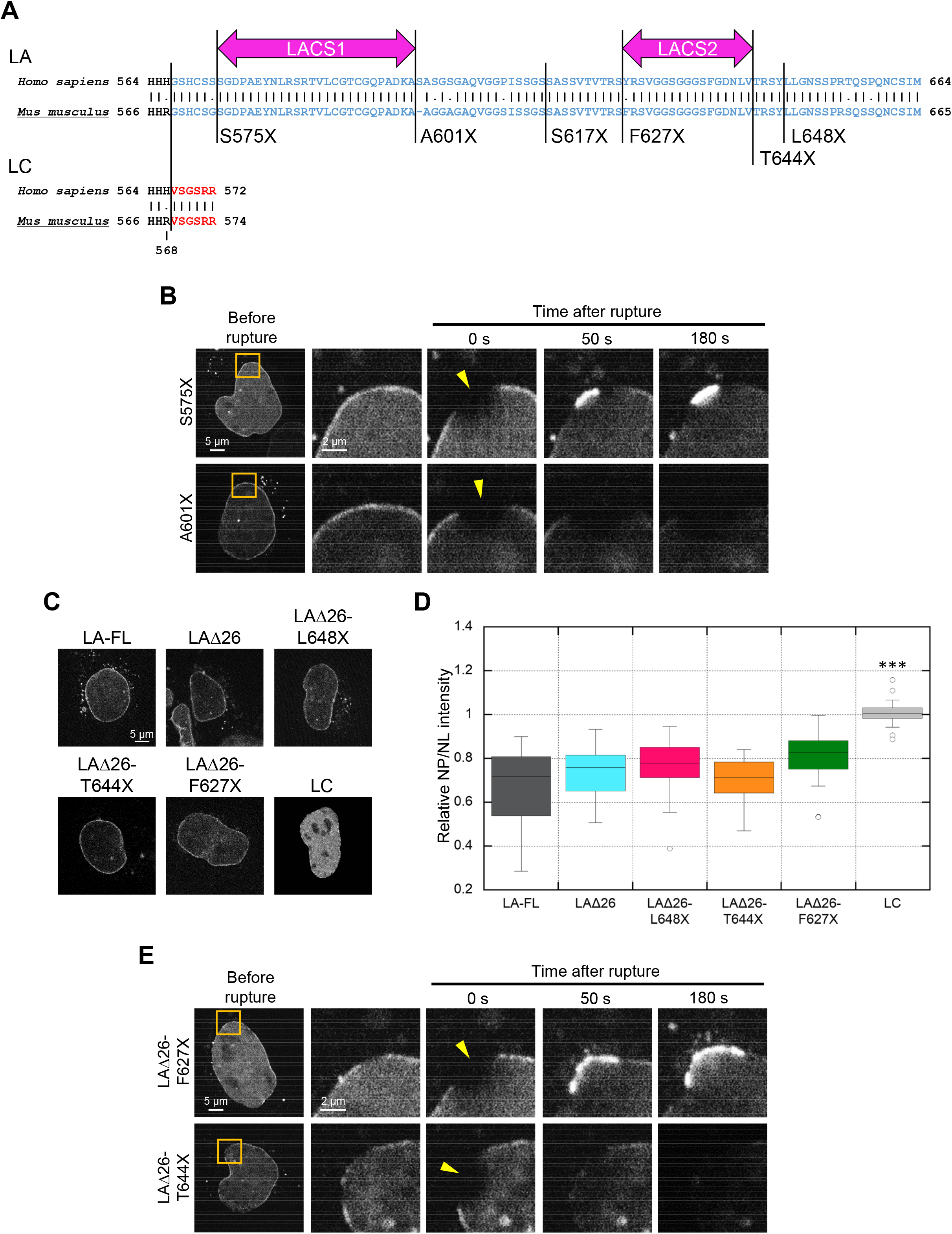
Accumulation kinetics of LA and PG mutants at the rupture sites in LA/C-KO MEFs. **(A)** Amino acid sequences of LA-(blue) or LC-specific (red) regions from *Homo sapiens* and *Mus musculus*. The amino acid homology is indicated as bars and a gap is introduced between residue A600 and A601 in *Mus musculus* to maximize the homology. **(B)** Dynamics of the mEmerald-LA-S575X and A601X in response to NE rupture in LA/C-KO MEFs. **(C)** Representative images of mEmerald-LA-full-length (FL), LA-FL without LACS1 (LAΔ26), LAΔ26-L648X, LAΔ26-T644X, LAΔ26-F627X, and mEmerald-LC in LA/C-KO MEFs transfected for 2 d. Bar: 5 μm. **(D)** Fluorescence intensities in the NP to the NL ratio of mEmerald-LA mutants and mEmerald-LC, as control, were measured (n = 14-20 cells from two independent experiments; ***, P < 0.001 by a Games–Howell test). (**C** and **D**) LA-FL and LC are reproduction of Fig. 5 B and C. **(E)** Dynamics of the mEmerald-LAΔ26-F627X and LAΔ26-T644X in response to NE rupture in LA/C-KO MEFs. **(B** and **E)** Magnified views of the indicated areas by orange boxes are shown (the second to fifth columns). A 2-μm diameter spot was laser-microirradiated to induce NE rupture (yellow arrowheads). Bars: 5 μm (the first column) and 2 μm (the second to fifth columns).

## References

1. J. M. Zullo et al., DNA sequence-dependent compartmentalization and silencing of chromatin at the nuclear lamina. Cell 149, 1474–1487 (2012).

2. M. Crisp et al., Coupling of the nucleus and cytoplasm: role of the LINC complex. J Cell Biol 172, 41–53 (2006).

3. G. K. Voeltz, M. M. Rolls, T. A. Rapoport, Structural organization of the endoplasmic reticulum. EMBO Rep 3, 944–950 (2002).

4. P. L. Paine, L. C. Moore, S. B. Horowitz, Nuclear envelope permeability. Nature 254, 109–114 (1975).

5. R. Foisner, L. Gerace, Integral Membrane-Proteins of the Nuclear-Envelope Interact with Lamins and Chromosomes, and Binding Is Modulated by Mitotic Phosphorylation. Cell 73, 1267–1279 (1993).

6. L. Gerace, A. Blum, G. Blobel, Immunocytochemical localization of the major polypeptides of the nuclear pore complex-lamina fraction. Interphase and mitotic distribution. J Cell Biol 79, 546–566 (1978).

7. L. Gerace, G. Blobel, The nuclear envelope lamina is reversibly depolymerized during mitosis. Cell 19, 277–287 (1980).

8. F. Lin, H. J. Worman, Structural organization of the human gene encoding nuclear lamin A and nuclear lamin C. J Biol Chem 268, 16321–16326 (1993).

9. T. H. Höger, K. Zatloukal, I. Waizenegger, G. Krohne, Characterization of a second highly conserved B-type lamin present in cells previously thought to contain only a single B-type lamin. Chromosoma 99, 379–390 (1990).

10. G. Biamonti et al., The gene for a novel human lamin maps at a highly transcribed locus of chromosome 19 which replicates at the onset of S-phase. Mol Cell Biol 12, 3499–3506 (1992).

11. F. Lin, H. J. Worman, Structural organization of the human gene (LMNB1) encoding nuclear lamin B1. Genomics 27, 230–236 (1995).

12. H. Maeno, K. Sugimoto, N. Nakajima, Genomic structure of the mouse gene (Lmnb1) encoding nuclear lamin B1. Genomics 30, 342–346 (1995).

13. T. Shimi et al., Structural organization of nuclear lamins A, C, B1, and B2 revealed by superresolution microscopy. Mol Biol Cell 26, 4075-4086 (2015).

14. Y. Turgay et al., The molecular architecture of lamins in somatic cells. Nature 543, 261–264 (2017).

15. T. Shimi et al., The A- and B-type nuclear lamin networks: microdomains involved in chromatin organization and transcription. Genes Dev 22, 3409–3421 (2008).

16. M. Kittisopikul et al., Computational analyses reveal spatial relationships between nuclear pore complexes and specific lamins. J Cell Biol 220 (2021).

17. R. J. Lutz, M. A. Trujillo, K. S. Denham, L. Wenger, M. Sinensky, Nucleoplasmic localization of prelamin A: implications for prenylation-dependent lamin A assembly into the nuclear lamina. Proc Natl Acad Sci U S A 89, 3000–3004 (1992).

18. M. Sinensky et al., The processing pathway of prelamin A. J Cell Sci 107 (Pt 1), 61–67 (1994).

19. V. L. Boyartchuk, M. N. Ashby, J. Rine, Modulation of Ras and a-factor function by carboxyl-terminal proteolysis. Science 275, 1796–1800 (1997).

20. A. M. Winter-Vann, P. J. Casey, Post-prenylation-processing enzymes as new targets in oncogenesis. Nat Rev Cancer 5, 405–412 (2005).

21. G. K. Leung et al., Biochemical studies of Zmpste24-deficient mice. J Biol Chem 276, 29051–29058 (2001).

22. D. P. Corrigan et al., Prelamin A endoproteolytic processing in vitro by recombinant Zmpste24. Biochem J 387, 129–138 (2005).

23. C. M. Denais et al., Nuclear envelope rupture and repair during cancer cell migration. Science 352, 353–358 (2016).

24. M. Raab et al., ESCRT III repairs nuclear envelope ruptures during cell migration to limit DNA damage and cell death. Science 352, 359–362 (2016).

25. C. T. Halfmann et al., Repair of nuclear ruptures requires barrier-to-autointegration factor. J Cell Biol 218, 2136–2149 (2019).

26. J. D. Vargas, E. M. Hatch, D. J. Anderson, M. W. Hetzer, Transient nuclear envelope rupturing during interphase in human cancer cells. Nucleus 3, 88–100 (2012).

27. J. Robijns et al., In silico synchronization reveals regulators of nuclear ruptures in lamin A/C deficient model cells. Sci Rep 6, 30325 (2016).

28. N. Y. Chen et al., Fibroblasts lacking nuclear lamins do not have nuclear blebs or protrusions but nevertheless have frequent nuclear membrane ruptures. Proc Natl Acad Sci U S A 115, 10100–10105 (2018).

29. N. Y. Chen et al., An absence of lamin B1 in migrating neurons causes nuclear membrane ruptures and cell death. Proc Natl Acad Sci U S A 116, 25870–25879 (2019).

30. Y. Kono et al., Nucleoplasmic lamin C rapidly accumulates at sites of nuclear envelope rupture with BAF and cGAS. J Cell Biol 221 (2022).

31. H. J. Worman, G. Bonne, “Laminopathies”: a wide spectrum of human diseases. Exp Cell Res 313, 2121–2133 (2007).

32. W. H. De Vos et al., Repetitive disruptions of the nuclear envelope invoke temporary loss of cellular compartmentalization in laminopathies. Hum Mol Genet 20, 4175–4186 (2011).

33. A. J. Earle et al., Mutant lamins cause nuclear envelope rupture and DNA damage in skeletal muscle cells. Nat Mater 19, 464–473 (2020).

34. P. H. Kim, et al., Nuclear membrane ruptures underlie the vascular pathology in a mouse model of Hutchinson-Gilford progeria syndrome. JCI Insight 6 (2021).

35. A. De Sandre-Giovannoli et al., Lamin a truncation in Hutchinson-Gilford progeria. Science 300, 2055 (2003).

36. M. Eriksson et al., Recurrent de novo point mutations in lamin A cause Hutchinson-Gilford progeria syndrome. Nature 423, 293–298 (2003).

37. A. M. Young, A. L. Gunn, E. M. Hatch, BAF facilitates interphase nuclear membrane repair through recruitment of nuclear transmembrane proteins. Mol Biol Cell 31, 1551–1560 (2020).

38. R. M. Sears, K. J. Roux, Mechanisms of A-Type Lamin Targeting to Nuclear Ruptures Are Disrupted in LMNA- and BANF1-Associated Progerias. Cells 11 (2022).

39. O. A. Zhironkina et al., Mechanisms of nuclear lamina growth in interphase. Histochem Cell Biol 145, 419–432 (2016).

40. S. Cho et al., Progerin phosphorylation in interphase is lower and less mechanosensitive than lamin-A,C in iPS-derived mesenchymal stem cells. Nucleus 9, 230–245 (2018).

41. I. C. Lopez-Mejia et al., A conserved splicing mechanism of the LMNA gene controls premature aging. Hum Mol Genet 20, 4540–4555 (2011).

42. F. G. Osorio et al., Splicing-directed therapy in a new mouse model of human accelerated aging. Sci Transl Med 3, 106ra107 (2011).

43. I. L. Ivanovska, M. P. Tobin, T. Bai, L. J. Dooling, D. E. Discher, Small lipid droplets are rigid enough to indent a nucleus, dilute the lamina, and cause rupture. J Cell Biol 222 (2023).

44. M. Zwerger et al., Altering lamina assembly reveals lamina-dependent and -independent functions for A-type lamins. J Cell Sci 128, 3607–3620 (2015).

45. V. Kochin et al., Interphase phosphorylation of lamin A. J Cell Sci 127, 2683–2696 (2014).

46. T. Shimi, C. G. Pack, R. D. Goldman, Analyses of the Dynamic Properties of Nuclear Lamins by Fluorescence Recovery After Photobleaching (FRAP) and Fluorescence Correlation Spectroscopy (FCS). Methods Mol Biol 1411, 99–111 (2016).

47. D. Holtz, R. A. Tanaka, J. Hartwig, F. McKeon, The CaaX motif of lamin A functions in conjunction with the nuclear localization signal to target assembly to the nuclear envelope. Cell 59, 969–977 (1989).

48. M. H. Gelb et al., Therapeutic intervention based on protein prenylation and associated modifications. Nat Chem Biol 2, 518–528 (2006).

49. R. Lee et al., Genetic studies on the functional relevance of the protein prenyltransferases in skin keratinocytes. Hum Mol Genet 19, 1603–1617 (2010).

50. S. Y. Chang et al., Inhibitors of protein geranylgeranyltransferase-I lead to prelamin A accumulation in cells by inhibiting ZMPSTE24. J Lipid Res 53, 1176–1182 (2012).

51. H. Yao et al., Targeting RAS-converting enzyme 1 overcomes senescence and improves progeria-like phenotypes of ZMPSTE24 deficiency. Aging Cell 19, e13200 (2020).

52. X. Chen et al., A small-molecule ICMT inhibitor delays senescence of Hutchinson-Gilford progeria syndrome cells. Elife 10 (2021).

53. K. N. Dahl et al., Distinct structural and mechanical properties of the nuclear lamina in Hutchinson-Gilford progeria syndrome. Proc Natl Acad Sci U S A 103, 10271–10276 (2006).

54. A. Quigley et al., The structural basis of ZMPSTE24-dependent laminopathies. Science 339, 1604–1607 (2013).

55. K. R. Yu et al., MicroRNA-141-3p plays a role in human mesenchymal stem cell aging by directly targeting ZMPSTE24. J Cell Sci 126, 5422–5431 (2013).

56. L. G. Fong et al., Heterozygosity for Lmna deficiency eliminates the progeria-like phenotypes in Zmpste24-deficient mice. Proc Natl Acad Sci U S A 101, 18111–18116 (2004).

57. A. Casasola et al., Prelamin A processing, accumulation and distribution in normal cells and laminopathy disorders. Nucleus 7, 84–102 (2016).

58. R. D. Goldman et al., Accumulation of mutant lamin A causes progressive changes in nuclear architecture in Hutchinson-Gilford progeria syndrome. Proc Natl Acad Sci U S A 101, 8963–8968 (2004).

59. T. Shimi et al., The role of nuclear lamin B1 in cell proliferation and senescence. Genes Dev 25, 2579–2593 (2011).

60. A. Ivanov et al., Lysosome-mediated processing of chromatin in senescence. J Cell Biol 202, 129–143 (2013).

61. Z. Dou et al., Autophagy mediates degradation of nuclear lamina. Nature 527, 105–109 (2015).

62. P. Scaffidi, T. Misteli, Reversal of the cellular phenotype in the premature aging disease Hutchinson-Gilford progeria syndrome. Nat Med 11, 440–445 (2005).

63. O. Santiago-Fernández et al., Development of a CRISPR/Cas9-based therapy for Hutchinson-Gilford progeria syndrome. Nat Med 25, 423–426 (2019).

64. J. L. V. Broers et al., Dynamics of the nuclear lamina as monitored by GFP-tagged A-type lamins. J Cell Sci 112 (Pt 20), 3463–3475 (1999).

65. T. Haraguchi et al., Nuclear localization of barrier-to-autointegration factor is correlated with progression of S phase in human cells. J Cell Sci 120, 1967–1977 (2007).

66. T. Haraguchi et al., Live cell imaging and electron microscopy reveal dynamic processes of BAF-directed nuclear envelope assembly. J Cell Sci 121, 2540–2554 (2008).

67. V. Stierlé et al., The carboxyl-terminal region common to lamins A and C contains a DNA binding domain. Biochemistry 42, 4819–4828 (2003).

68. A. C. Schibler, P. Jevtic, G. Pegoraro, D. L. Levy, T. Misteli, Identification of epigenetic modulators as determinants of nuclear size and shape. Elife 12, 10.1101/2022.1105.1128.493845 (2023).

69. K. Ikegami, S. Secchia, O. Almakki, J. D. Lieb, I. P. Moskowitz, Phosphorylated Lamin A/C in the Nuclear Interior Binds Active Enhancers Associated with Abnormal Transcription in Progeria. Dev Cell 52, 699–713 e611 (2020).

70. V. T. Chu et al., Increasing the efficiency of homology-directed repair for CRISPR-Cas9-induced precise gene editing in mammalian cells. Nat Biotechnol 33, 543–548 (2015).

71. Y. Guo, Y. Kim, T. Shimi, R. D. Goldman, Y. Zheng, Concentration-dependent lamin assembly and its roles in the localization of other nuclear proteins. Mol Biol Cell 25, 1287–1297 (2014).

72. M. Roblek et al., Monoclonal antibodies specific for disease-associated point-mutants: lamin A/C R453W and R482W. PLoS One 5, e10604 (2010).

73. B. Marcos-Ramiro et al., Isoprenylcysteine Carboxylmethyltransferase-Based Therapy for Hutchinson-Gilford Progeria Syndrome. ACS Cent Sci 7, 1300–1310 (2021).

74. A. Sánchez-López et al., Cardiovascular Progerin Suppression and Lamin A Restoration Rescue Hutchinson-Gilford Progeria Syndrome. Circulation 144, 1777–1794 (2021).

75. H. Park, S. S. Han, Y. Sako, C. G. Pack, Dynamic and unique nucleolar microenvironment revealed by fluorescence correlation spectroscopy. FASEB J 29, 837–848 (2015).

76. Y. Kanda, Investigation of the freely available easy-to-use software ‘EZR’ for medical statistics. Bone Marrow Transplant 48, 452–458 (2013).

77. D. Bates, M. Mächler, B. M. Bolker, S. C. Walker, Fitting Linear Mixed-Effects Models Using lme4. Journal of Statistical Software 67, 1–48 (2015).

78. A. Kuznetsova, P. B. Brockhoff, R. H. B. Christensen, lmerTest Package: Tests in Linear Mixed Effects Models. Journal of Statistical Software 82, 1–26 (2017).

79. M. Mirdita et al., ColabFold: making protein folding accessible to all. Nat Methods 19, 679–682 (2022).

